# Niche-inspired synthetic matrices for epithelial organoid culture

**DOI:** 10.1101/806919

**Authors:** Victor Hernandez-Gordillo, Timothy Kassis, Arinola Lampejo, GiHun Choi, Mario E. Gamboa, Juan S. Gnecco, David T. Breault, Rebecca Carrier, Linda G. Griffith

**Affiliations:** Center for Gynepathology Research and Biological Engineering Department, Massachusetts Institute of Technology, 77 Massachusetts AV Cambridge MA, USA 02138.; Deparment of Pediatrics, Harvard Medical School, 300 Longwood Avenue, Boston, MA 02115.; Department of Chemical Engineering, Northeastern University, 208 Lake Hall, Boston, MA, USA, 02115.

## Abstract

Epithelial organoids are now an important tool in fields ranging from regenerative medicine to drug discovery. Organoid culture requires Matrigel, a complex, tumor-derived, extracellular matrix. An alternative completely synthetic matrix could improve culture reproducibility, clarify mechanistic phenomena, and enable applications involving human implantation. Here, we designed synthetic matrices with tunable biomolecular and biophysical properties that allowed us to identify critical gel parameters in organoid formation. Inspired by known epithelial integrin expression in the proliferative niche of the human intestine, we identified an α2β1 integrin-binding peptide as a critical component of the synthetic matrix that supports human duodenal colon and endometrial organoid propagation. We show that organoids emerge from single cells, retain their proliferative capacity, are functionally responsive to basolateral stimulation and have correct apicobasal polarity upon induction of differentiation. The local biophysical presentation of the cues, rather than bulk mechanical properties, appears to be the dominant parameter governing epithelial cell proliferation and organoid formation in the synthetic matrix.

## Introduction

Epithelial organoids are three-dimensional (3D) structures with microarchitecture, cellular composition and functions, similar to their native tissues^1^. They are used to model tissue homeostasis and pathophysiological processes and as *in vitro* models for drug development^1–3^. However, their full potential has been hindered by the requirement for propagation in Matrigel, a complex, tumor-derived extracellular matrix (ECM) containing a plethora of growth factors (GFs) that are variable from lot-to-lot and exert unknown effects on organoid phenotype^4–7^. Envisioned applications involving re-implantation of patient-derived organoids require Good Manufacturing Practices (GMP) that exclude tumor-derived material^8, 9^. Hence, there are many forces driving the development of synthetic matrices in which all exogenous biological cues are known and defined.

Towards this goal, Gjorevski et al., developed a synthetic poly(ethylene) glycol (PEG) gel modified with a fibronectin (FN)-derived RGD peptide or laminin (LMN)-derived peptides that supported mouse intestinal organoid growth but not differentiation. Differentiation was achieved in a semisynthetic matrix made with PEG and full-length LMN^10^. Human organoids survived in the PEG-RGD gel, although expansion, differentiation, and other features were not reported. Mouse organoid growth in the PEG gel appear to depend on the gel’s bulk mechanical properties: stiff (∼1.3 kPa) for organoid formation and soft (∼300 Pa) for differentiation^10^. The gel’s bulk mechanical properties were modulated by processes that simultaneously also affect other gel properties, such as ligand accessibility, clustering, and nanomechanical responses to cell-generated forces after ligand binding^11–19^. The observation that organoids grow significantly better in Matrigel or in hydrogels made with low concentrations of purified LMN than in PEG-RGD or collagen gels^10, 20–23^ suggests that specific cell-matrix affinity interactions might play a critical role in organoid formation and differentiation. Untangling the relative contributions of bulk mechanical properties from other gel biophysical and biomolecular properties on cell proliferation and organoid formation is challenging.

Inspired by integrins expression in the native tissue, here we create synthetic matrices with tunable biophysical and biomolecular properties that support organoid formation and differentiation across multiple donors and tissue types. The local biophysical presentation and the identity of the integrin-binding motif, rather than bulk biomechanical properties or matrix degradation, appear to be the dominant variables governing epithelial cell proliferation and organoid formation in the synthetic matrix.

### Niche-inspired synthetic ECM

Intestinal stem cells (ISCs) at the bottom of the crypt express α2β1 and α5β1 integrins ^24–29^. Consequently, ligands for these receptors are attractive components of synthetic ECM. We created a modular synthetic ECM by combining 8-arm vinyl sulfone-activated PEG macromers partially modified with integrin-binding peptides and matrix-binding peptides together with peptide crosslinkers containing a matrix-metalloproteinase (MMP)-sensitive degradation site. In our design, we focused on three variables that influence integrin-ligand interactions: a) identity of the integrin-binding ligand b) biophysical properties of ligand presentation and c) matrix biomechanical properties at local and macroscale (Fig 1a). We also included cell-mediated matrix remodeling (via proteases and matrix deposition) and dynamic softening, via Sortase degradation, as additional variables.

**Fig 1.**
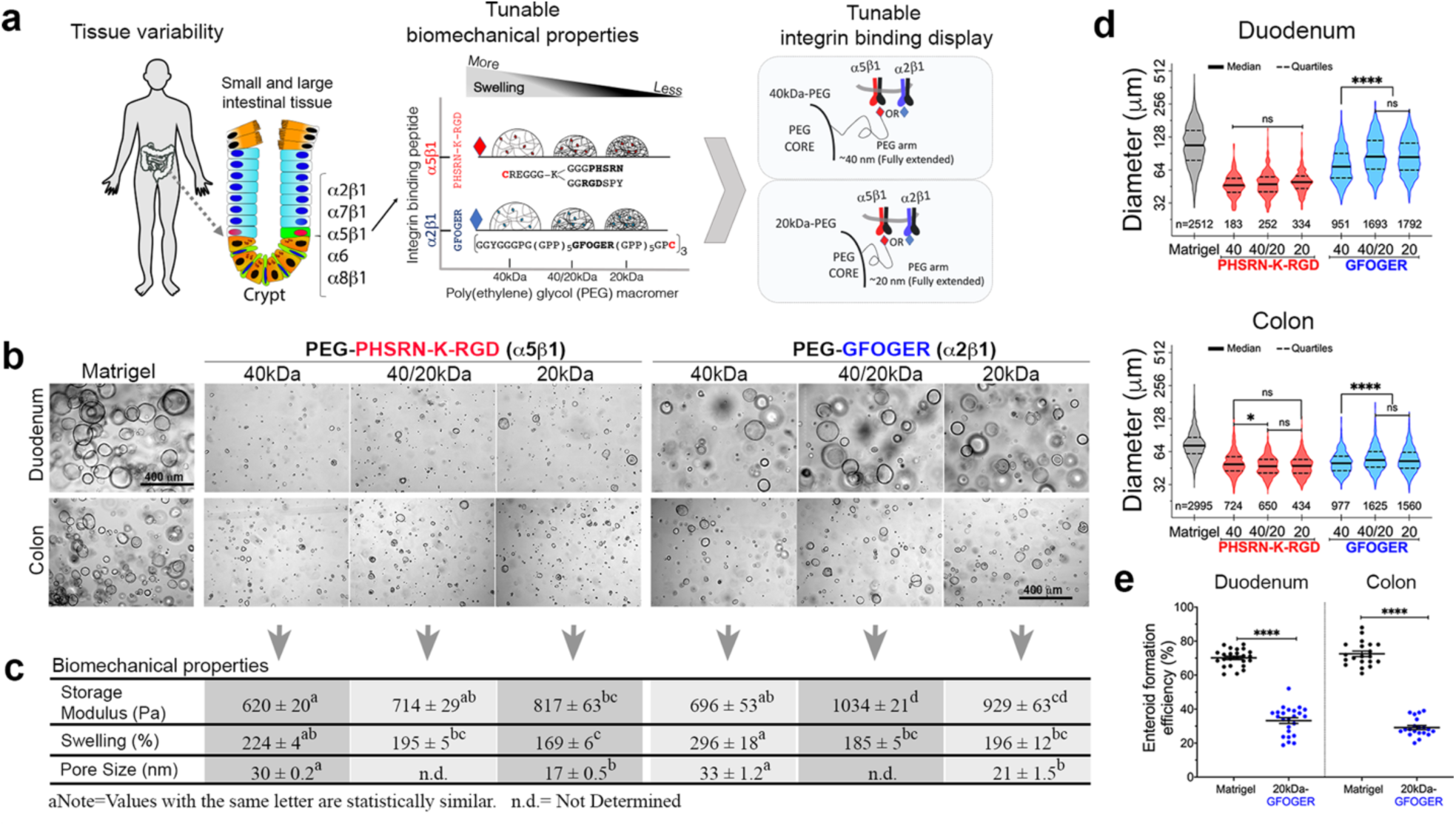
Synthetic matrices support enteroid formation from single cells. **a,** Niche-inspired synthetic matrix design. **b,** Representative images of six-day old duodenal and colon enteroids emerging in the synthetic matrices. Seven donors (six human duodenal, one colon, and one mouse colon) were evaluated (Extended Data Fig 2). Scale bar: 400 μm. **c,** Biomechanical properties of the synthetic matrices. Storage modulus data (n=6 gels). Swelling and theoretical pore size data (n=12 gels). **d,** Enteroid diameter of duodenal (n=3 experiments) and colon (n=2 experiments) donors. The number of enteroids measured is depicted under each violin plot. Matrigel diameters were significantly bigger (P<0.0001). **e**, Enteroid formation efficiency of duodenunal and colon donors from two independent experiments. Each symbol represents a single hydrogel with the mean and SEM.

As PEG chains are highly flexible in aqueous solution, they can be tailored in length to modulate ligand accessibility^11–19^ and in combination with crosslinking density, also modulate local and bulk mechanical properties^30–32^. In our design, the arm PEG lengths were modulated to either ∼20 nm or ∼40 nm (when fully extended from the central polymer core) by using 8-arm PEG macromers of 20kDa or 40kDa, respectively (Fig 1a). To parse the role of the α2β1 and α5β1 integrins on cell survival and organoid formation, we created matrices using single, or mixed (1:1, w/v) macromers with a fixed crosslinking density. To engage the α5β1 integrin, we functionalized the PEG macromers with an FN-derived peptide that contains the ***RGD*** motif and the ***PHSRN*** synergy site (PEG-***PHSRN-K-RGD***)^11, 33, 34^. Likewise, to engage the α2β1 integrin, we used a collagen I-derived peptide harboring the ***GFOGER*** sequence (PEG-***GFOGER***)^35^.

We also considered that initial integrin engagement is only a first step, with possible subsequent interactions mediated by matrix that cells produce and assemble locally^36, 37^. Hence, we included peptides with affinity for FN, collagen IV (C-IV), and LMN (FN-binder and BM-binder, respectively)^38–41^. Lastly, we considered that organoid growth might require matrix degradation, thus we used a protease-sensitive crosslinker concatenated with a site susceptible to cleavage with exogenously-added transpeptidase SrtA enzyme to enable cell-mediated matrix remodeling and recovery of intact organoids, respectively^38^.

Six synthetic matrix formulations (40kDa-***PHSRN-K-RGD***, 40kDa-***GFOGER***, 20kDa-***PHSRN-K-RGD***, 20kDa-***GFOGER***, 40/20kDa-***PHSRN-K-RGD***, 40/20kDa-***GFOGER***) that varied in bulk mechanical properties but contained similar crosslinker density and equal nominal concentration of integrin-, BM- and FN-binding peptides, were made. In the PEG-***PHSRN-K-RGD*** matrices, the **RGD** and **PHSRN** sites are presented in a branched configuration to mimic the biophysical presentation found in native FN^11, 42^ whereas in the PEG-***GFOGER***, three cell-binding cues are presented as a triple helix, thus mimicking the biophysical presentation of collagen fibers (Fig 1a)^35^.

### Synthetic ECM supports enteroid growth from different donors

Donor-to-donor and species variability is known to affect organoid growth^43, 44^. Thus, to test the synthetic matrices, we selected seven human donors (six duodenal and one colon) that varied in sex, age and pathological state, and one mouse colon donor. Dissociated single cells were embedded in the synthetic matrices and then cultured in expansion medium (EM). To facilitate the identification of a suitable matrix for organoid culture, we used the organoid diameter as a proxy for cell proliferation and organoid growth (Extended Data Fig 1)^45^.

Overall, organoids emerging in both Matrigel and in the synthetic matrices had a wide range of size distributions, but all were spherical with a single layer of thin epithelial cells. We considered this phenotype as undifferentiated and potentially stem-enriched organoids (Fig 1b, Extended Data Fig 2a). We designated these spherical organoids as “enteroids” and reserved the term “organoid” for enteroids that underwent differentiation and adopted a thick columnar epithelial cell layer.

The synthetic matrices supported enteroid formation from human and mouse donors with various degree of success (Fig 1b,d, Extended Data Fig 2). Compared to Matrigel, human enteroids in the synthetic matrices were smaller. Interestingly, human colon enteroids grew slower, in Matrigel and the synthetic matrices, compared to the duodenal enteroids. In contrast, mouse colon enteroids grew faster than human colon enteroids in Matrigel or the synthetic matrices. In some matrix conditions mouse colon enteroids reached sizes similar to mouse colon enteroids in Matrigel, (Extended Data Fig 2b).

From the two sets of synthetic matrices, the PEG-***GFOGER*** (α2β1) series supported enteroid formation, across all donors, to a greater extent than the PEG-***PHSRN-K-RGD*** (α5β1) series, even though the encapsulated cells expressed α2, α5, and β1 integrins (Extended Data Fig 3a). The ability of the PEG-***GFOGER*** series to support enteroid formation is not due to bulk mechanical differences, as the 20kDa-***PHSRN-K-RGD*** and the 20kDa-***GFOGER*** matrices have similar mechanical properties, yet we observed significant differences in enteroid diameters (Fig 1c-d, Extended Data Fig 2). We also noted that the ***GFOGER*** bioactivity is context dependent, as matrices with relatively high swelling, lower storage modulus, and longer PEG arms, failed to promote robust enteroid formation (Compare 40kDa-***GFOGER*** vs 20kDa-***GFOGER***). Matrices made with the 1:1 (w/v) mixed macromers (40/20kDa-***GFOGER***) rescued the ***GFOGER*** bioactivity. This is more likely because ∼70% of PEG arms in the 40/20kDa-***GFOGER*** matrix correspond to the 20kDa-PEG macromer, which results in a matrix of similar biomechanical properties as the 20kDa-***GFOGER*** (Fig 1b-d). This phenomenon of context-dependent success of enteroid formation was not observed in the PEG-***PHSRN-K-RGD*** series, in part because enteroid emergence was relatively poor.

Collectively, the data shows that intestinal cell proliferation and enteroid formation occurs in a narrow set of matrix mechanical properties in human enteroids and in a broader range in mouse enteroids. Notably, our results highlight the significance of using a variety of donors and tissue types, as matrices that supported mouse enteroids failed with human donors. Across the human donors, we also noticed donor-to-donor variability on enteroid diameter and size distribution in the synthetic matrices that mirrored the donor-to-donor variability observed in Matrigel. Compared to Matrigel, the synthetic ECM supported enteroid formation efficiency of 47% (duodenal) and 40% (colon) (Fig 1e). These differences could be due to the number of residual growth factors and cytokines in Matrigel that are absent in our synthetic ECM.

### Enteroid growth depends on GFOGER/α2β1

Modulation of the ligand’s properties in the 20kDa-***GFOGER*** matrix revealed that enteroid formation depends on the α2β1 integrin binding to the ***GFOGER*** peptide (Fig 2a) in a dose-dependent manner (Fig 2b-c). An inactive form of the ***GFOGER*** peptide (***GFOG<u>D</u>R***)^35^ caused an increase in the number of dead cells and failed to promote enteroid formation (Fig 2a and c). Thus, engaging α2β1 is not only essential for enteroid formation but also needed to maintain a population of cells that do not form enteroids, yet remain viable.

**Fig 2.**
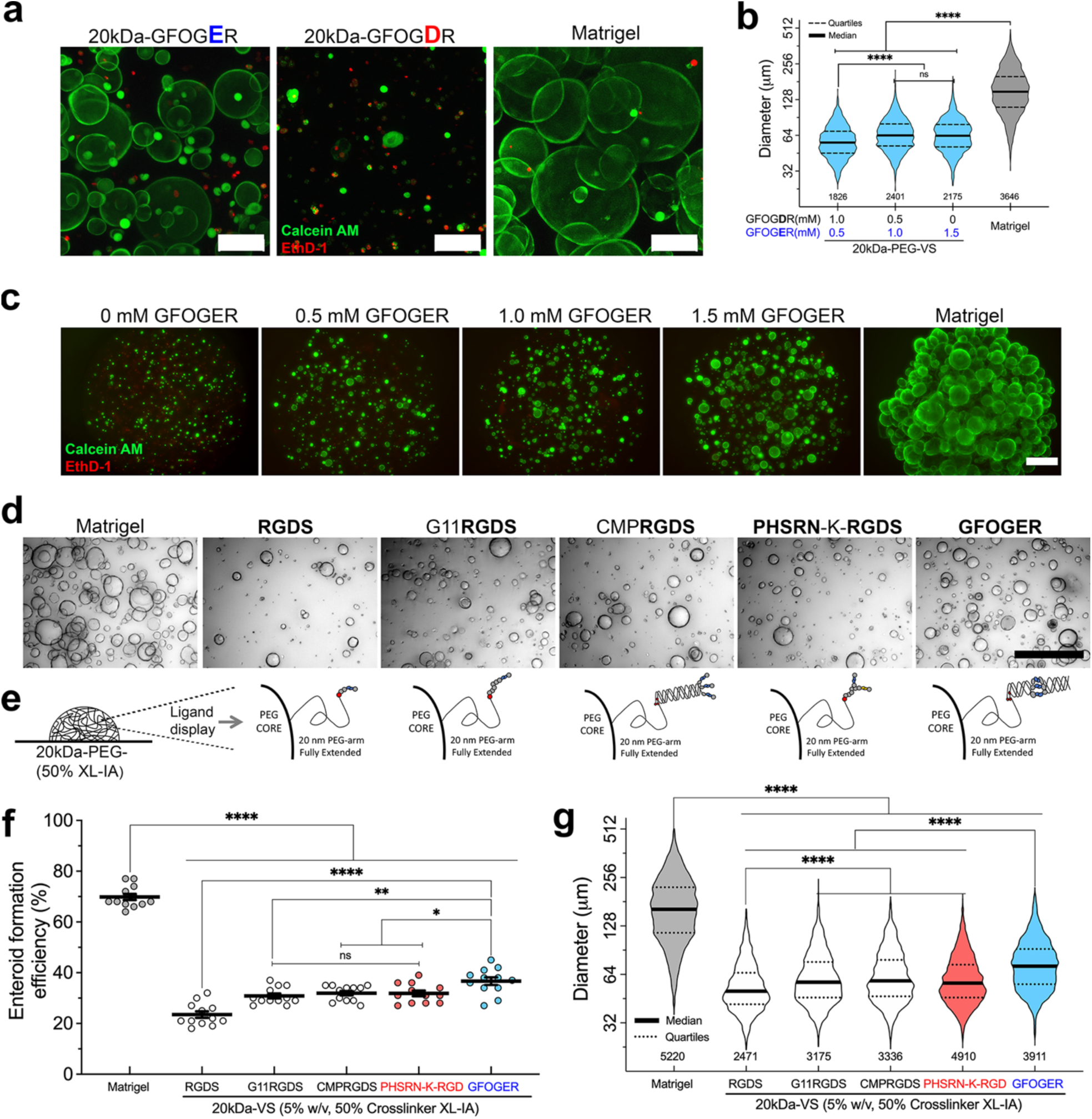
Robust enteroid growth depends on the α2β1 integrin binding peptide, GFOGER. **a**, An aminoacidic point-mutation in GFOGER (E◊D) abolishes enteroid formation. Live-dead 3D projections of enteroids. **b-c**, GFOGER dose response effect on **b**, enteroid diameter and **c**, viability. **d-f**, Effect of the α5β1 ligand presentation on enteroid formation efficiency and diameter. **d**, Enteroids in matrices with various ligand presentation in **e**. **f**, Enteroid formation efficiency from two experiments. Each symbol represents a single hydrogel with the mean and SEM. **g**, Enteroid diameters from two independent experiments. The number of enteroids measured per matrix condition in **b** and **g** is depicted under each violin plot. Scale bar in **a**: 100 μm, **c-d**: 500 μm. Six-day old enteroids were used/analyzed in **a**-**d** and **g**.

Changes in the identity and concentration of the crosslinker also affected enteroid formation. Enteroids emerged from single cells encapsulated in a non-degradable synthetic ECM, but by 6 days were smaller and less abundant compared to enteroids in matrices made with a cell-degradable crosslinker (Extended Data Fig 3b). This suggests that the matrix’s microarchitecture can support initial cell proliferation to establish enteroids, but later stages in growth might require simultaneous matrix degradation. Crosslinker densities of 45 and 50% at 1.5 mM nominal concentration of ***GFOGER*** appears the most favorable environment for enteroid growth. Lower crosslinker densities caused greater hydrogel swelling and reduced enteroid growth, possible due to changes in ligand distribution combined with a dilution effect of the ***GFOGER*** concentration within the matrix (Extended Data Fig 3c).

Because ligand presentation and clustering affect integrin binding, we wondered if the higher ***GFOGER*** bioactivity compared to ***PHSRN-K-RGDS*** was due to the triple-helical nature of the peptide that extends and clusters the ligand away from the PEG polymer core compared to the branched and shorter configuration of the ***PHSRN-K-RGDS***. Longer and clustered integrin binding peptides have been shown to increase accessibility and integrin binding which results in higher cell proliferation^46–48^. Thus, we hypothesized that increasing the α5β1 ligand accessibility and clustering, while maintaining the bulk mechanical properties constant, would result in increased enteroid formation. We synthesized three peptides harboring the α5β1 ligand (Arg-Gly-Asp,Ser, RGDS); a short linear peptide (***RGDS***), a longer extended ligand (***G11RGDS***) and a longer and clustered **RGDS** ligand (***CMPRGDS***)^47^ (Fig 2d-e). Although we saw an increase in enteroid formation efficiency and diameters with the extended (***G11RGDS***) and clustered (***CMPRGDS***) peptides compared to linear ***RGDS***, they were significantly smaller were less efficient in enteroid formation than the ***GFOGER*** matrix (Fig 2f-g).

Collectively, the previous data point to a mechanism for enteroid formation that appears to be dependent on the ***GFOGER***-α2β1 integrin interaction, as extending and clustering the a5β1 (**RGDS**) ligand failed to promote robust enteroid growth. Further, ***GFOGER*** presentation appears to be favored by shorter (∼20 nm) compared to longer (∼40 nm) PEG arms, possibly due to a favorable nanomechanical response when the cells pull on the ligand^11, 13, 17, 19^.

### Enteroid characterization in the synthetic ECM

To gain more insight into the growth of human enteroids in the synthetic ECM, we followed the formation of enteroids from single cells (Fig 3a, Extended Data Fig 4a). Time-lapse live imaging from days 1 to 2 and 2 to 3 after encapsulation revealed faster growth in Matrigel compared to the synthetic ECM (**Supplementary videos**). A clear lumen was visible by day three in Matrigel and by day four in the synthetic ECM (Fig 3a). By day six, cells that failed to form enteroids were still viable (Fig 3b). Viable single cells that did not form enteroids were observed in various human donors (Extended Data Fig 4b). Time-lapse live cell imaging from day 4 to 6 revealed that enteroids in Matrigel and the synthetic ECM grew in a manner reminiscent of fluid pressure oscillations^49, 50^. Time-lapse live cell imaging also captured epithelial cells being extruded into the lumen in Matrigel and the synthetic ECM, suggesting the establishment of a functional and polarized epithelial layer (see red arrows in Fig 3d and **supplementary videos**). Actin staining further confirmed correct enteroid polarization across multiple donors (Extended Data Fig 4c).

**Fig 3.**
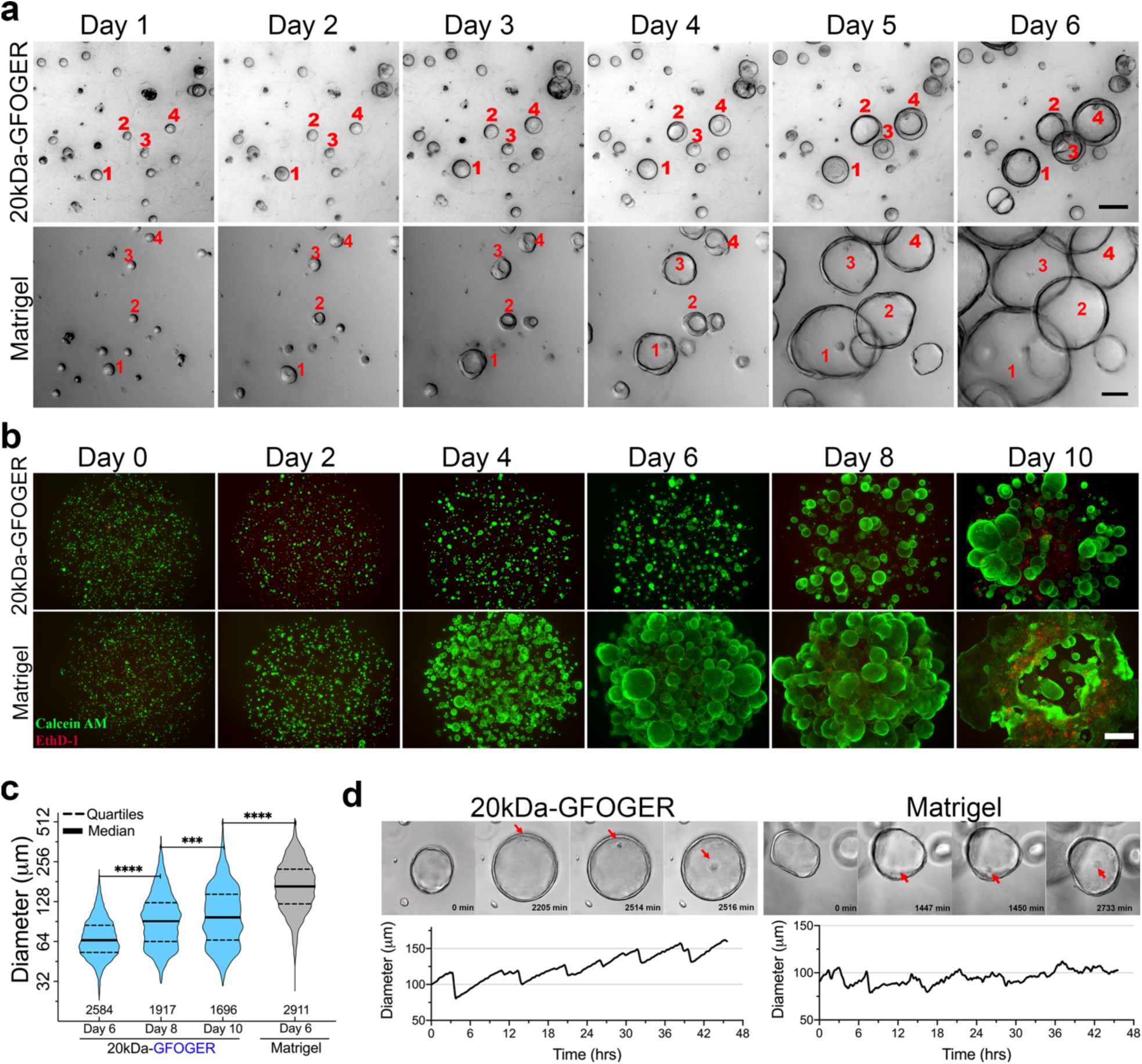
Single cell proliferation and enteroid formation in the synthetic ECM are similar to those in Matrigel. **a**, Time-course imaging of enteroids emerging from single cells. Scale bar: 50 μm. **b**, Live-dead 3D projections of enteroids over ten days. Scale bar: 500 μm. **c**, Time-course enteroid diameter (n=2 experiments). The number of enteroids measured is depicted under each violin plot. **d**, Time-lapsed imaging analysis of changes in diameter of a single enteroid from days 4 to 6. Red arrows point to single cells from the epithelial layer being extruded into the lumen.

As enteroids in the synthetic ECM, on average, had a delayed growth, we extended our organoid diameter analysis for up to ten days. Although enteroids in the synthetic ECM continued to grow (Fig 3b-c), the size distribution of the overall population at day 10 did not reach that of the population in Matrigel at day 6. Interestingly, enteroids in Matrigel at days 8 and 10 started to fuse and created a mass of interconnected cells at the bottom of the well accompanied by significant cell death, possibly due to Matrigel degradation and dissolution. We did not observe such a phenomenon in the synthetic ECM, which is an important feature for long-term culture and emerging application of organoids.

Properly polarized and functional organoids can be used to study adaptive epithelial responses^51^. To test if enteroids in the synthetic ECM were functionally responsive to basolateral stimulation, we treated six-day old enteroids with prostaglandin E2 (PGE2) and forskolin (FKL). PGE2 and FKL have been shown to induce rapid morphological changes characterized by an increase in organoid diameter^51, 52^. Upon PGE2 and FKL treatment, enteroids in the synthetic ECM increased in diameter, similar to the response we observed with enteroids in Matrigel (Extended Data Fig 6d-e, Supplementary videos). This response suggests that the synthetic ECM not only supports functionally responsive enteroids but is also flexible enough to accommodate a rapid increase in enteroid size that is more likely independent of matrix degradation.

### Enteroids in the synthetic ECM retain their proliferative capacity

Epithelial organoids can be cultured continuously due to an undifferentiated stem cell population that, when re-embedded in Matrigel, gives rise to new organoids^53^. To determine if a population of undifferentiated and proliferative cells existed in enteroids emerging in the synthetic ECM, we performed quantitative PCR (qPCR) for five genes associated with active stem cells, six genes associated with quiescent stem cells (+4 position) and nine genes associated with progenitor cells ^54, 55^. Overall, in the two duodenal donors tested, we observed similar levels of gene expression of active and quiescent stem and progenitor genes in enteroids emerging in the synthetic ECM compared to Matrigel, except for SMOC2, MMP7, and ATOH1 genes that showed more variability (Fig 4a and Extended Data Fig 5a).

**Fig 4.**
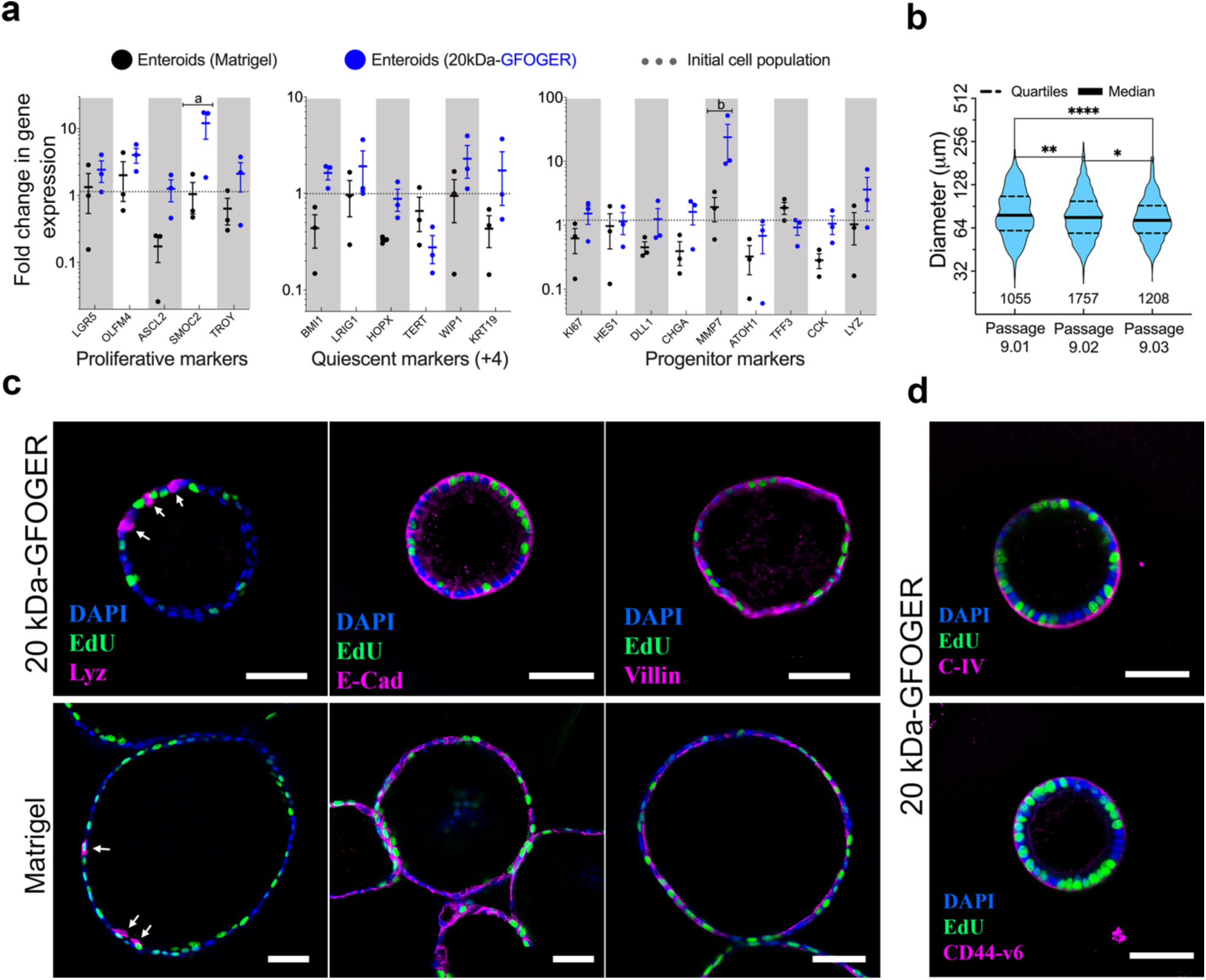
Enteroids in the synthetic ECM show diverse cellular composition. **a**, Fold change in gene expression of enteroids in the synthetic ECM or Matrigel. Data from three independent experiments. **b**, Quantification of enteroid diameters from three successive passages in the synthetic ECM. Data from a single experiment with 12 replicates in each passage. The number of enteroids measured is depicted under each violin plot. **c-d**, Immunostaining analysis of six-day old enteroids in the synthetic ECM and Matrigel showing proliferative cells (Edu), Paneth cells (Lyz), epithelial markers (E-cad and Villin) and basolateral markers (C-IV and CD44-v6). Scale bar: 50 μm. Six-day old enteroids were used in all panels.

As enteroids in the synthetic matrix appear to retain a similar gene expression profile to enteroids in Matrigel, we sought to investigate if they contained a proliferative cell population capable of forming new enteroids when re-embedded in new synthetic ECM or Matrigel. Six-day old enteroids were collected from the synthetic ECM and then processed (see Methods) to generate single cells to re-embed into new synthetic ECM or Matrigel (Extended Data Fig 6a). We performed this procedure for three consecutive passages with two duodenal donors. Quantification of the total number of cells recovered after each passage in the synthetic ECM revealed a 1.5 to 2-fold increase in cell expansion relative to the initial number of encapsulated cells (Extended Data Fig 6b). As described before, we also noted a population of cells that failed to form enteroids but remained viable. This cell population was also included in the calculation of fold increase in Extended Data Fig 6b. Interestingly, when single cells from organoids grown in the synthetic ECM were embedded in Matrigel there was more than 14-fold increase in cell expansion as the majority of single cells formed enteroids. This might be due to the number of residual growth factors present in Matrigel that could induce a proliferative phenotype in otherwise quiescent cells^55, 56^. This observation furthers highlights the significance of developing a suitable matrix to uncover biological processes in epithelial organoids masked when using ill-defined Matrigel hydrogels. Quantification of enteroid diameters after each passage further revealed a wide range of sizes in the population, but similar to previous results, enteroids in Matrigel were, on average, bigger than those in the synthetic ECM (Fig 4b, Extended Data Fig 5b, 6c). The wide range in enteroid diameter suggests that enteroids recovered from the synthetic ECM, similar to those recovered from Matrigel, contain a population of cells that vary in their proliferative capacity.

To further characterize the duodenal enteroids in the synthetic ECM, we labeled them with 5-ethynyl-2’-deoxyuridine (EdU) to identify proliferative cells, then performed immunostaining to identify Paneth (Lyz) cells, as Paneth cells are part of the intestinal stem cell niche *in vivo*. We observed a number of EdU and lyz positive cells in enteroids grown in Matrigel and synthetic ECM in two donors tested. Further, enteroids also showed E-cadherin and villin, typical markers for intestinal epithelial cells Collagen IV and CD44-v6 (a hyaluronic acid receptor expressed at the bottom of the crypt)^57^ were also detected in enteroids growing in the synthetic ECM (Fig 4c-d and Extended Data Fig 5c).

### Enteroid differentiation in synthetic ECM

Stem-enriched enteroids in Matrigel undergo differentiation when switched to differentiation medium (DM) lacking Wnt3a, a ligand for the Frizzled receptor that enhances the activity of the LGR5 ligand, R-spondin1, in LGR5-positive ISCs^53^, or DM made with lower concentrations of L-WRN (Wnt3a, R-spondin1 and Noggin) conditioned medium (L-WRN-DM)^63^. Stem-enriched mouse enteroids grown in a synthetic ECM, in contrast, undergo rapid cell death upon switching to DM, due to a rapid loss of resident stem cells^10^. We hypothesized that to allow differentiation in a synthetic ECM we would need to preserve a pool of stem cells while allowing the emergence of differentiated cells. Thus, enteroids were differentiated using a lower concentration of L-WRN conditioned medium. Human duodenal and colon enteroids were grown for six days in EM (50% L-WRN) then switched to L-WRN-DM (25% L-WRN) or DM. Similar to previous results with mouse organoid differentiation in a synthetic ECM, removal of Wnt3a (DM) caused rapid cell death of human enteroids that was not observed in enteroids differentiated in Matrigel (Data not shown). In the presence of L-WRN-DM, human enteroids adopted diverse phenotypes with some enteroids showing typical hallmarks of differentiation (thick columnar cells and accumulation of apoptotic cells in the lumen), and in some instances adopting folded 3D structures (Fig 5a). Mouse intestinal enteroids in the synthetic ECM also showed hallmarks of differentiation (Extended Data Fig 9a-b). Immunostaining analysis of human and mouse organoids also revealed correct polarization as shown with a diverse array of apical (F-actin, NHE3, DPP4, villin) and basolateral markers (collagen IV, LMN, CD44v6). Proliferative cells (Ki67 or EdU), Paneth cells (Lyz), and Goblet cells (Muc2) were also observed in the differentiated duodenal and colon enteroids (Fig 5b-c, Extended Data Figure 7-8, 9c). In summary, the data shows that enteroids emerging in the synthetic ECM are composed of stem cells that can undergo differentiation to a similar extent to enteroids emerging in Matrigel.

**Fig 5.**
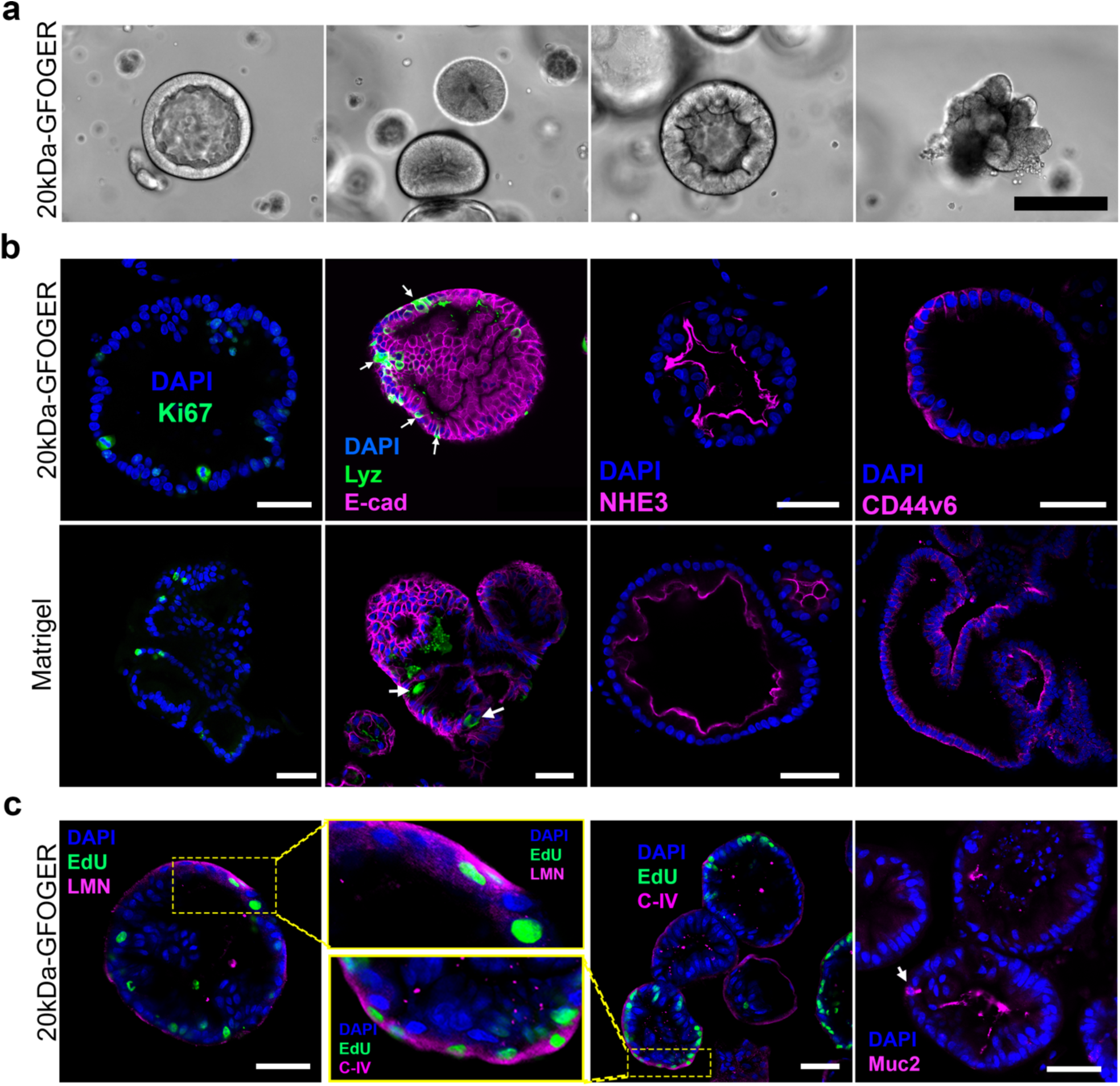
Enteroids in the synthetic ECM undergo differentiation. **a**, Bright-field images of 10-day old human enteroids growing in the synthetic ECM showing diverse phenotypes upon inducing differentiation. Scale bar: 200 μm. **b-c**, Immunostaining analysis of 10-day old organoids in the synthetic ECM and Matrigel showing proliferative cells (Ki67), Paneth cells (Lyz), Goblet cells (Muc2). Epithelial marker (E-cad), apical marker (NHE3) and basolateral markers (CD44v6, C-IV, and LMN). Scale bar: 500 μm.

### Endometrial organoids in synthetic ECM

We extended our study to include human endometrial organoids, as similar to intestinal organoids, they are composed of a single layer of epithelial cells surrounding a lumen. Interestingly, endometrial glands *in vivo* express the α2β1 and αVβ1 integrin receptors^59–61^. Thus, we selected a panel of synthetic matrices to engage the α2β1 (20kDa-***GFOGER***) or the αVβ1 (20kDa-***PHSRN-K-RGD***, 20kDa-***RGDS***, 20kDa-***G11RGDS*** and 20kDa-***CMPRGDS***) integrins. Single cells embedded in the synthetic matrices formed globular organoids with thick columnar cells surrounding an empty lumen. Similar to intestinal enteroids, endometrial organoids in Matrigel were bigger and more abundant than organoids in the synthetic matrices. Nonetheless, from all the synthetic matrices tested, the 20kDa-***GFOGER*** (α2β1) supported bigger and more abundant endometrial organoids (Fig 6a-b). Endometrial organoids, in the synthetic ECM, showed proliferative cells (EdU), epithelial (EpCAM) and apicobasal markers (actin, LMN) similar to those grown in Matrigel (Fig 6c-d).

**Figure 6.**
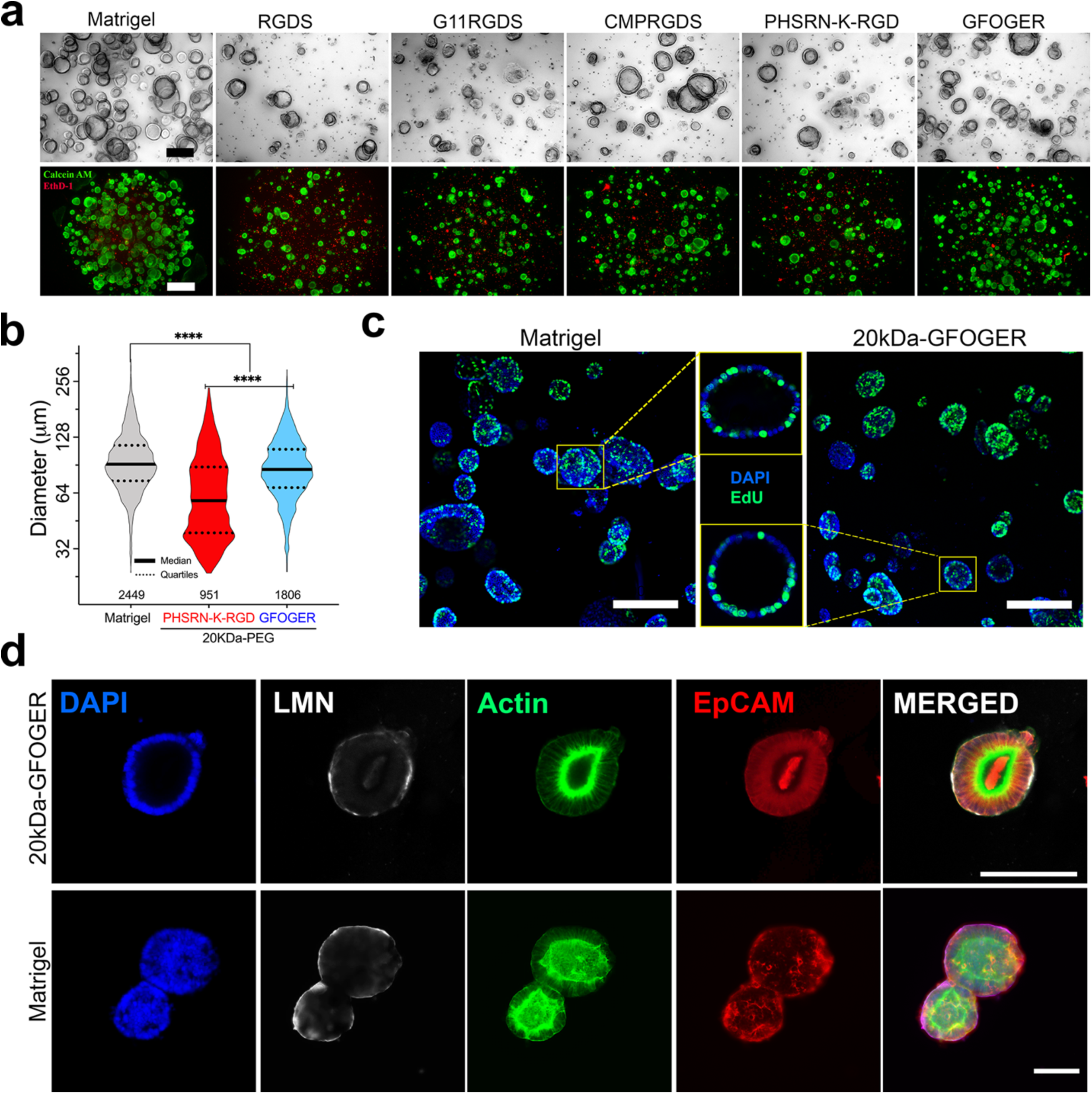
Human endometrial organoids can be cultured in a synthetic ECM. **a**, Top, Bright field images of endometrial organoids in the synthetic matrix and Matrigel, eight days after encapsulation. Bottom, live-dead 3D projections of endometrial organoids in the indicated matrix after eight days of culture. Scale bar top: 200 μm, bottom: 500 μm. **b**, Diameter of eigh-day old endometrial organoids (n=2 independent experiments). The number of organoids measured per matrix condition is depicted under each violin plot. **c-d**, Immunostaining analysis of organoids showing proliferative cells (Edu), apical markers (actin), epithelial marker (EpCAM), and basolateral marker (LMN). Scale bar in **c** and **d**, bottom panel = 200 μm, top panel in **d**: 100 μm.

## Conclusions

Designing a synthetic matrix that supports robust organoid formation and differentiation from a spectrum of different donor origins has been challenging due to differences between mouse and human organoids along with donor-to-donor variability. We report for the first time, to our knowledge, a fully synthetic matrix that support organoid formation from different tissue types and epithelial organs. Inspired by integrin expression in the native tissue, we found that the local biophysical presentation of an α2β1 integrin-binding peptide, rather than the bulk gel’s mechanical properties, appears to be the dominant variable governing epithelial cell proliferation and organoid formation. We found that enteroids emerging in the synthetic ECM are, on average, less numerous and showed delayed growth compared to Matrigel. This discrepancy could be attributed to: residual GFs in Matrigel that affect organoid biology in unknown ways; additional matrix components (e.g., laminins) that stimulate additional signaling pathways; the propensity of Matrigel to bind GFs present in the culture media and serve as a depot for later release; or any combination of these. Interestingly, single cells that do not progress to enteroids remain viable in the synthetic ECM for over a week. Thus, the synthetic matrix may allow interrogation and discovery of factors that impact organoid biology.

Although not fully explored here, the synthetic ECM offers additional benefits such as on-demand dissolution to recover intact cells and cell-secreted metabolites^38^ and incorporation of small peptide that sequester cell-secreted matrix for an in-depth investigation of cell-matrix deposition in tissue formation ^39^. We expect that these features will be of great utility to uncover complex and dynamic cell-cell communications in emerging stromal-epithelial co-culture systems^39^. We further envision that the synthetic matrix can also be used in microfabrication approaches aimed to replicate the intestinal topography^62, 63^. Such systems would also require cues to engage α2β1-expressing cells to preserve a proliferative compartment. In conclusion, the matrix developed here has the potential to be a “one size fits all” to multiple applications. To facilitate its adoption by the organoid and bioengineering community, we used commercially available and made-to-order reagents that require minimal manipulation and common laboratory equipment.

---

**Extended Data Fig 1.**
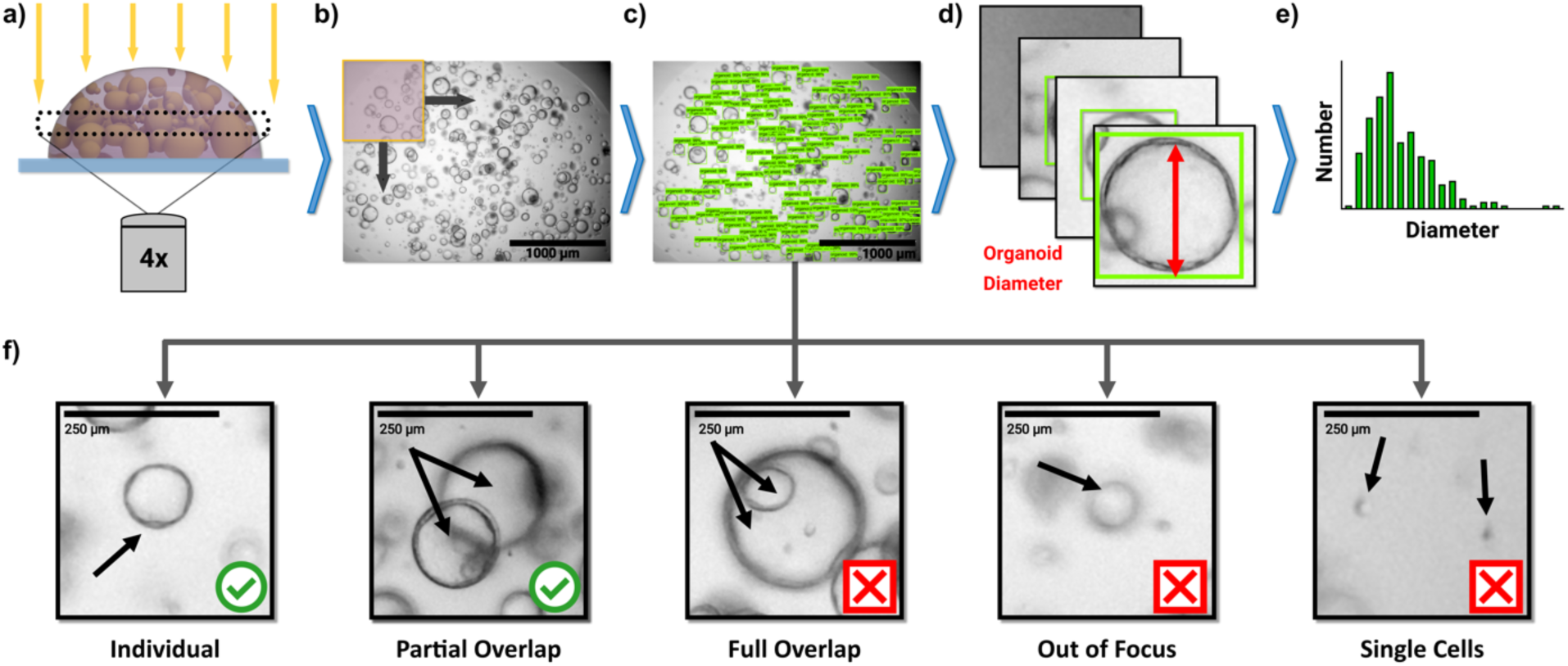
Enteroid diameter measurements using Orgaquant, a deep convolutional neural network analysis platform. **a,** A single plane, approximately in the middle of the matrix droplet, is imaged in bright-field (BF) using a 4x objective. **b**, The BF image is sectioned into smaller images using a sliding window of 450 pixels, with an overlap window of 200 pixels. The overlapping allows to capture organoids that otherwise would be at the edge of the slide window. The algorithm also eliminates redundant bounding boxes that might occur due to overlapping. **c**, A representative BF image showing bounding boxes around the enteroids measured. **d**, The diameter of the bounding boxes is used to calculate the diameter of the enteroids in focus, which are then presented as a distribution plot, as in **e**. **f**, The convolutional network was trained to ignore enteroids that have full overlap, are out of focus, or appear as single cells.

**Extended Data Fig 2.**
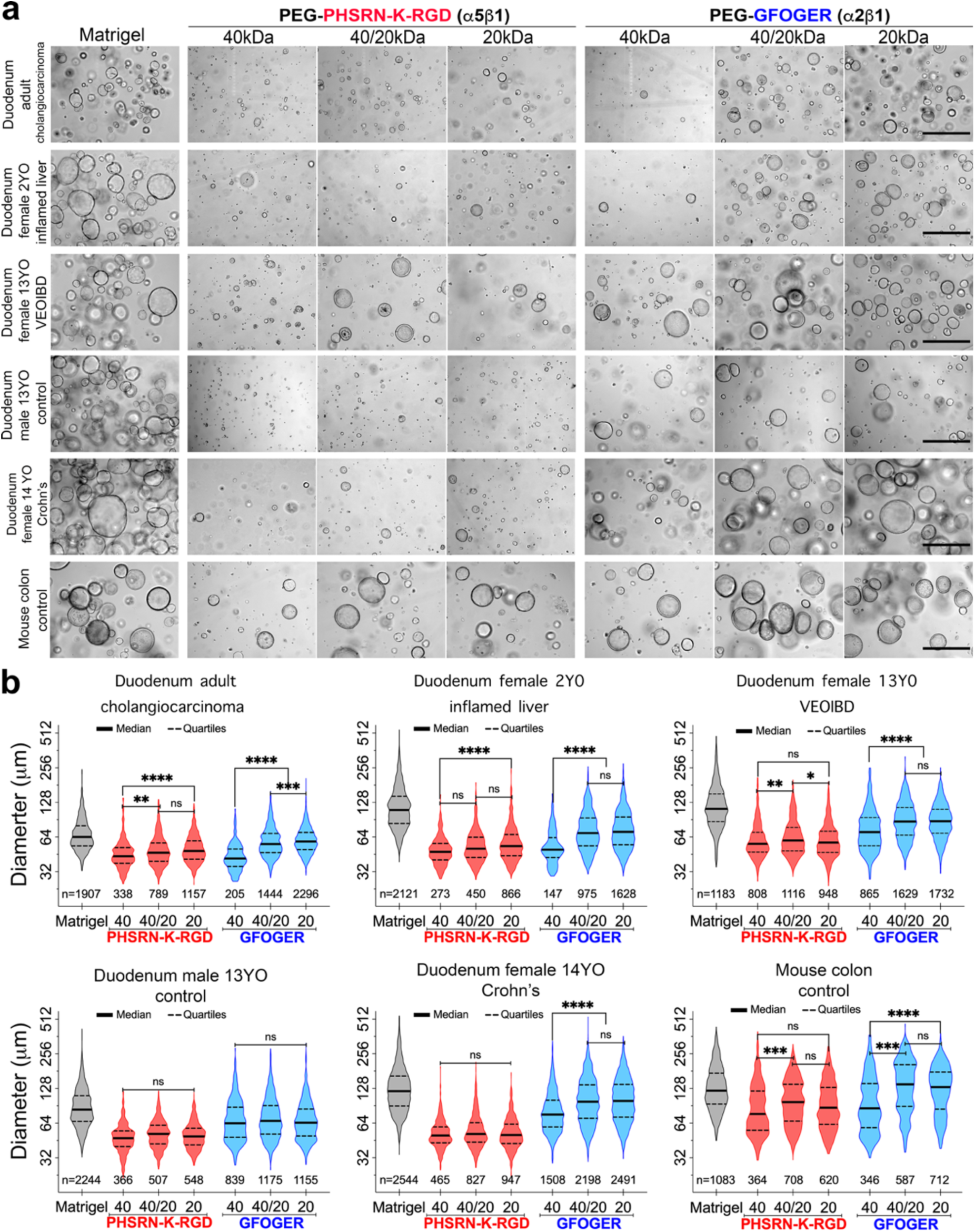
Synthetic matrices support enteroid formation from multiple donors and tissue types. **a**, Representative images of six-day (human) and four-day (mouse) enteroids growing in the synthetic matrices. Scale bars: 400 μm. **b,** Quantification of enteroid diameter from three independent experiments for human donors and two independent experiments for mouse donor. The number of enteroids measured per matrix condition and donor is depicted under each violin plot. Matrigel diameters were significantly bigger (P<0.0001).

**Extended Data Fig 3.**
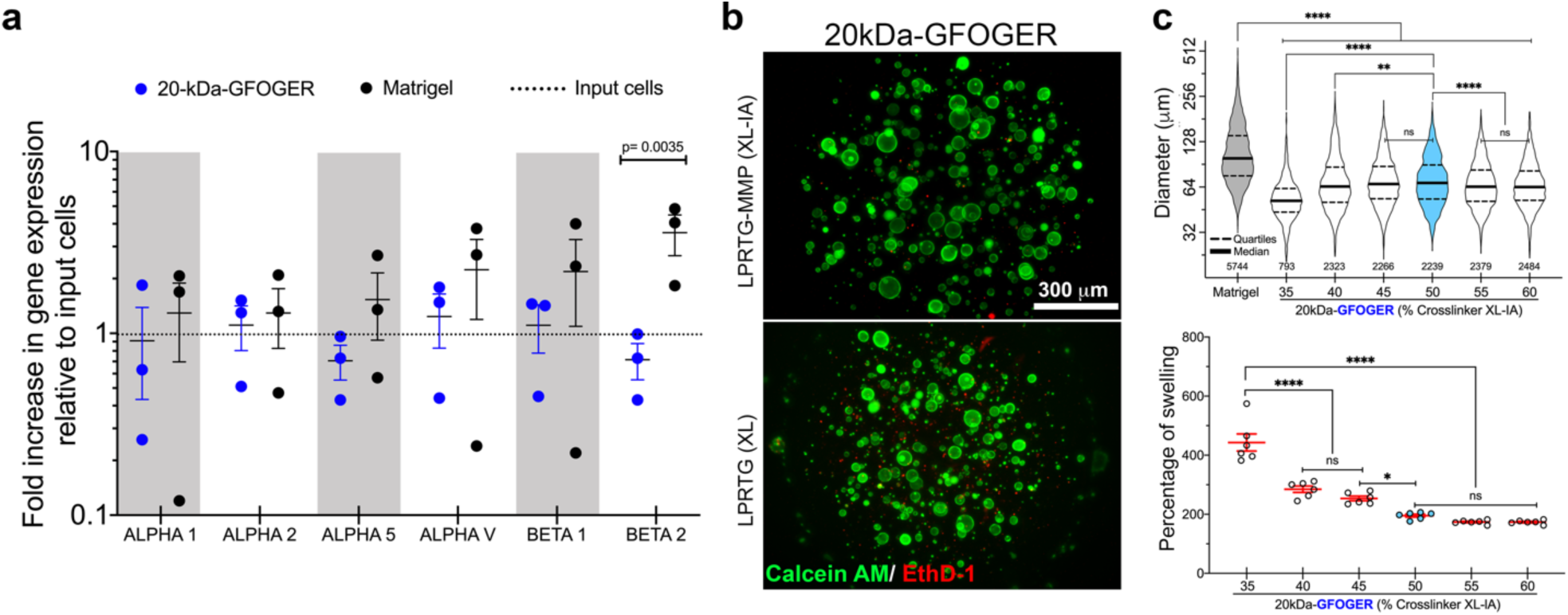
Integrin expression and matrix’s biomechanical properties affect enteroid formation. **a**, Integrin expression profile of six-day old human duodenal enteroids**. b,** Live/dead 3D projections of six-day old enteroids growing in matrices made with a degradable (LPRTG-MMP) and non-degradable (LPRTG) crosslinker. **c**, Effect of crosslinker density (LPRTG-MMP) on the enteroid diameter (top) and gel swelling property (bottom). Data from three independent experiments. The number of six-day old enteroids measured is depicted under each violin plot in the top panel. Bottom panel, each symbol represents a single hydrogel with the mean and SEM.

**Extended Data Fig 4.**
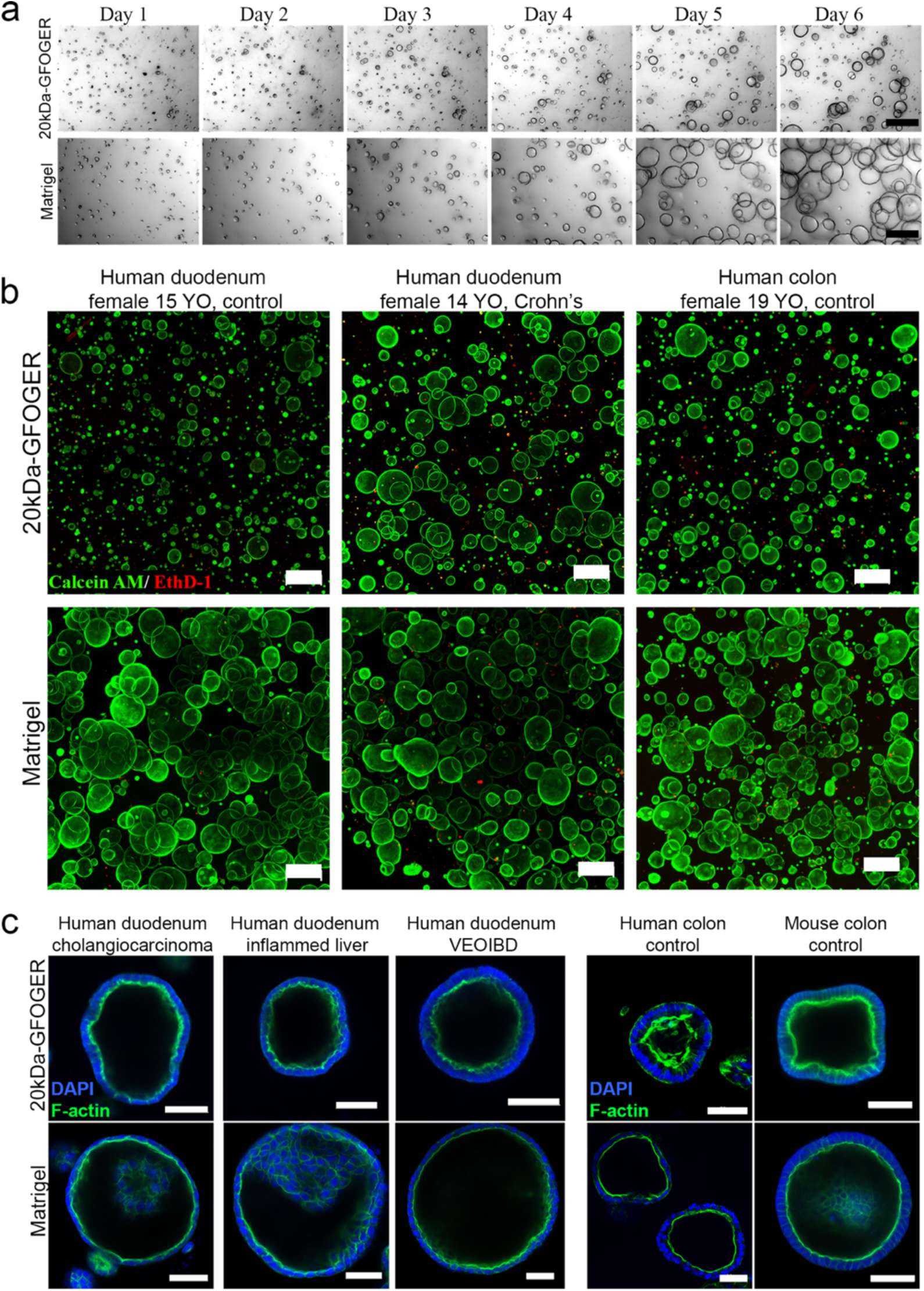
Single cell proliferation and enteroid formation in the synthetic ECM are similar to those in Matrigel. **a**, Time-course imaging of single cells forming enteroids in the 20kDa-GFOGER and Matrigel. Scale bar: 50 μm. **b**, Live-dead 3D projections of six-day (duodenal) and eight-day (colon) enteroids in the 20kDa-GFOGER and Matrigel. Scale bar: 500 μm. **c**, Human duodenal (left) and colon enteroids display well-defined and polarized lumens when grown in the synthetic ECM or Matrigel. Human duodenal enteroids were grown for six days, human colon for eight days, and mouse colon for four days.

**Extended Data Fig 5.**
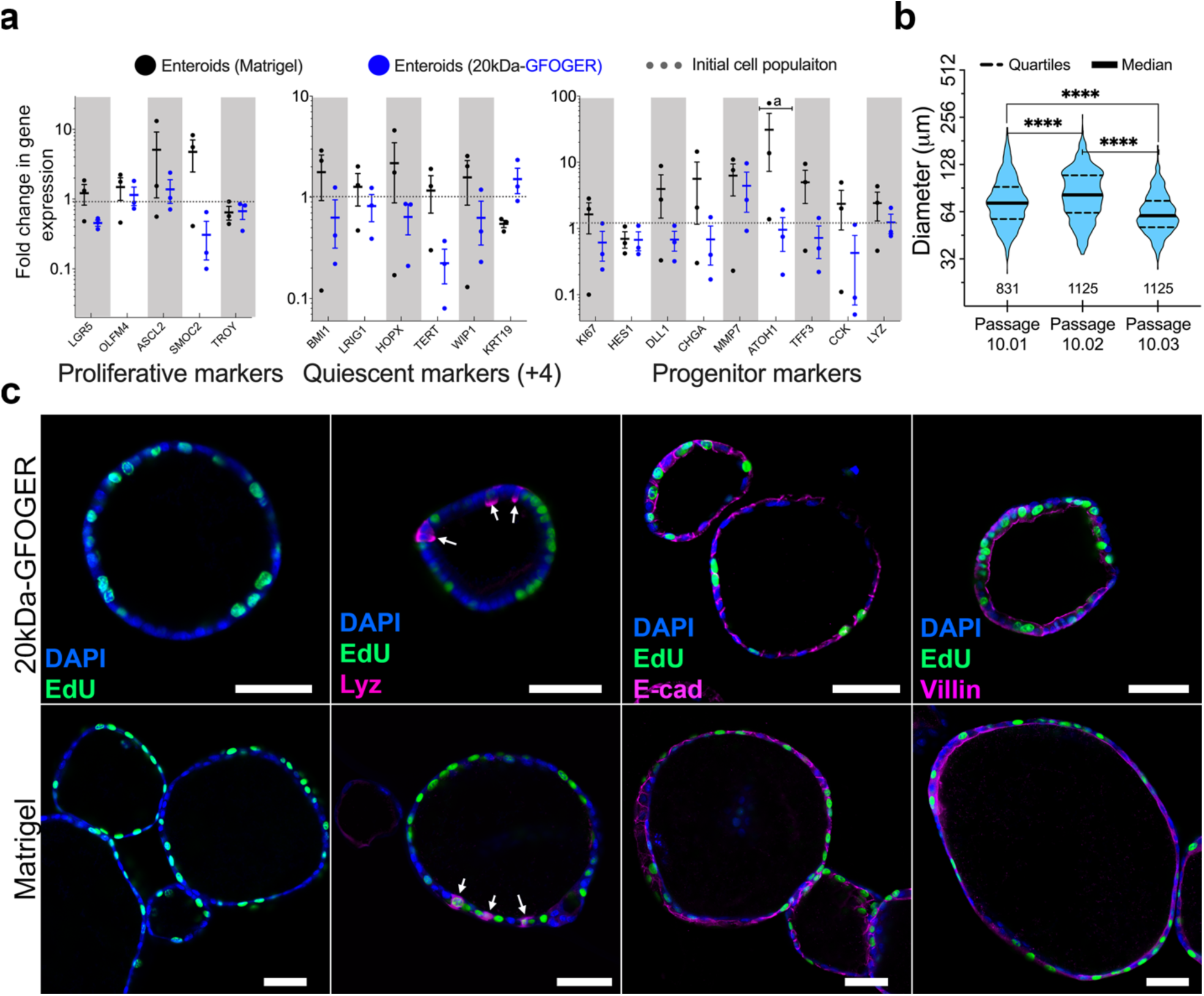
Human duodenal enteroids from a Crohn’s disease donor show correct cellular composition in the synthetic ECM. **a**, Fold change in gene expression of six-day old enteroids in the synthetic ECM and Matrigel relative to the input cells. Data from three independent experiments. **b**, Quantification of six-day old enteroid diameters from three successive passages in the synthetic ECM. Data from a single experiment with 12 replicates per consecutive passage. The number of enteroids measured is depicted under each violin plot. **c**, Immunostaining analysis of six-day old enteroids in the synthetic ECM and Matrigel showing proliferative cells (Edu), Paneth cells (Lyz) and epithelial markers (E-cad and Villin).

**Extended Data Fig 6.**
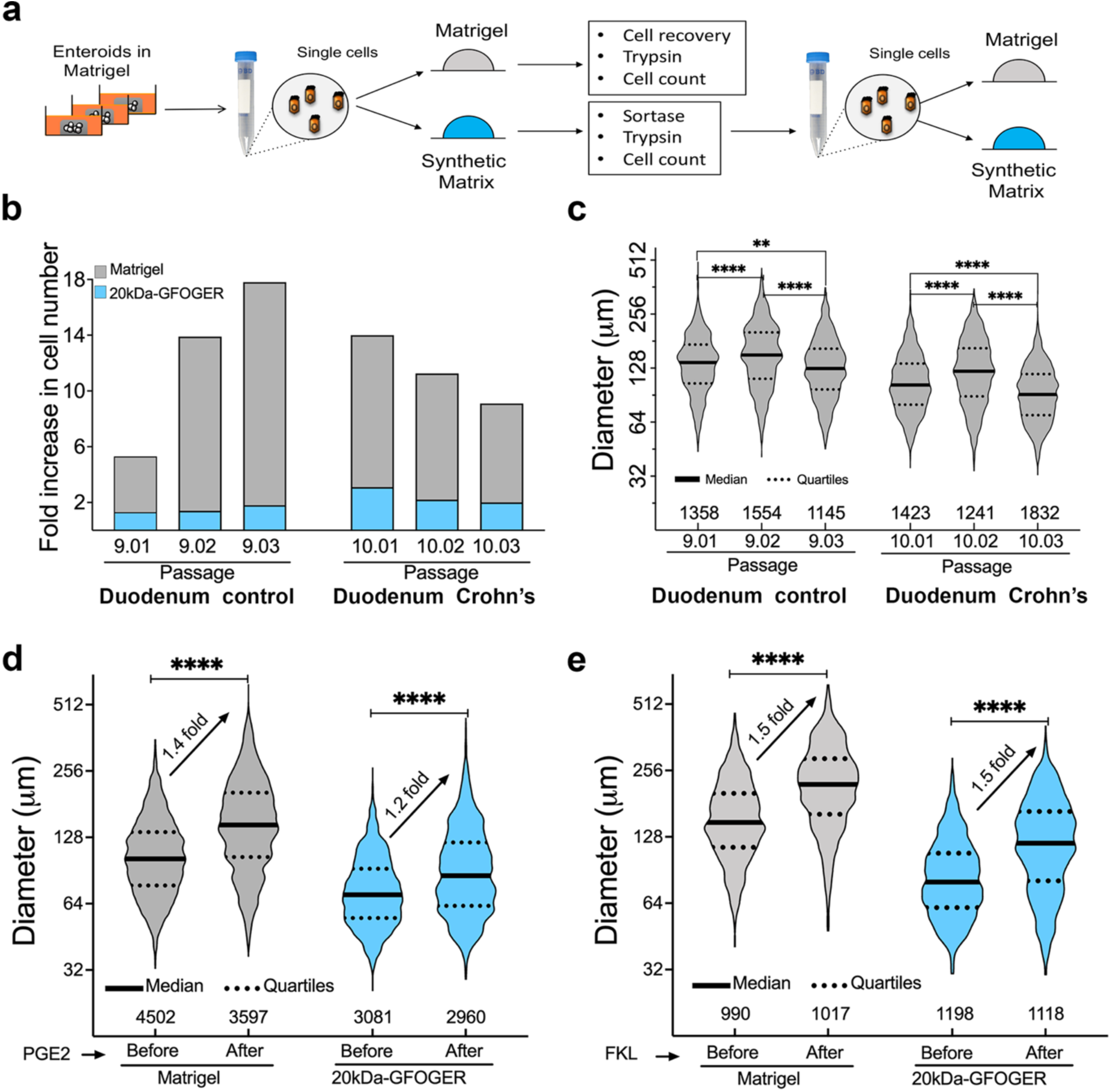
Enteroids grown in the synthetic ECM retain their proliferative capacity and respond to basal stimulation. **a**, Overall strategy to test the proliferative capacity of enteroids grown in the synthetic ECM. After Sortase-mediated dissolution, six-day old enteroids were digested and single cells re-embedded in new synthetic ECM or Matrigel. **b**, Fold increase in cell number after six days of culture in each passage from two human donors. Data represent three consecutive passages using enteroids collected from 24 to 48 individual hydrogels pooled. **c**, Diameter of six-day old enteroids in Matrigel emerging from enteroids previously grown in the synthetic ECM. Data from a single experiment with three consecutive passages. The number of enteroids measured is depicted under each violin plot. **d-e**, Diameter of enteroids before (day six) and after (day seven) stimulation with PGE2 (n=3) and forskolin (n=2). The number of enteroids measured is depicted under each violing plot.

**Extended Data Fig 7.**
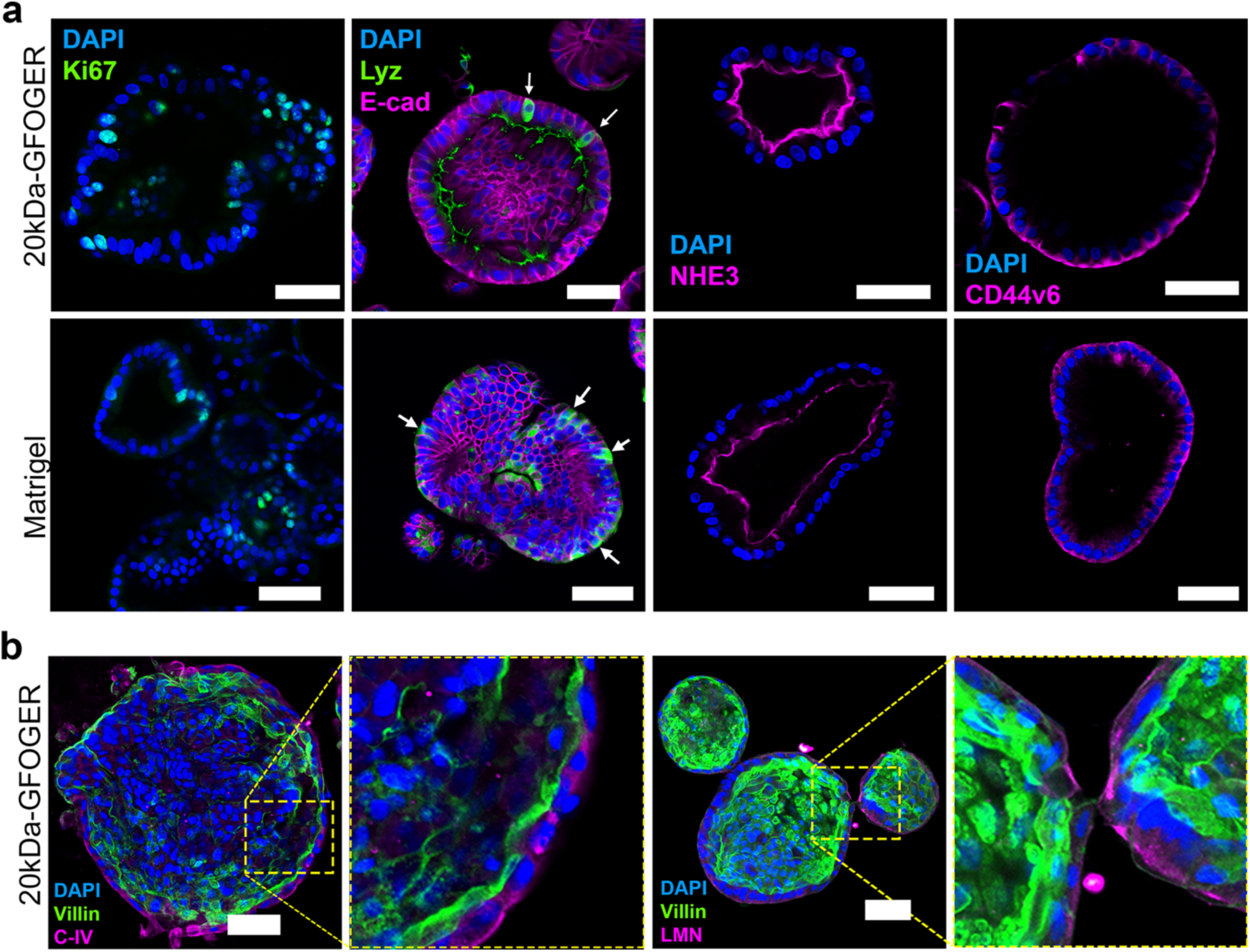
Human duodenal enteroids from a Crohn’s disease donor show appropriate cellular composition after differentiation. **a-b**, Immunostaining analysis of 10-day old organoids differentiated in the synthetic ECM and Matrigel showing proliferative cells (Ki67), Paneth cells (Lyz), epithelial marker (E-cad), apical markers (NHE3, Villin) and basolateral markers (CD44v6, C-IV, and LMN). Scale bar: 50 μm.

**Extended Data Fig 8.**
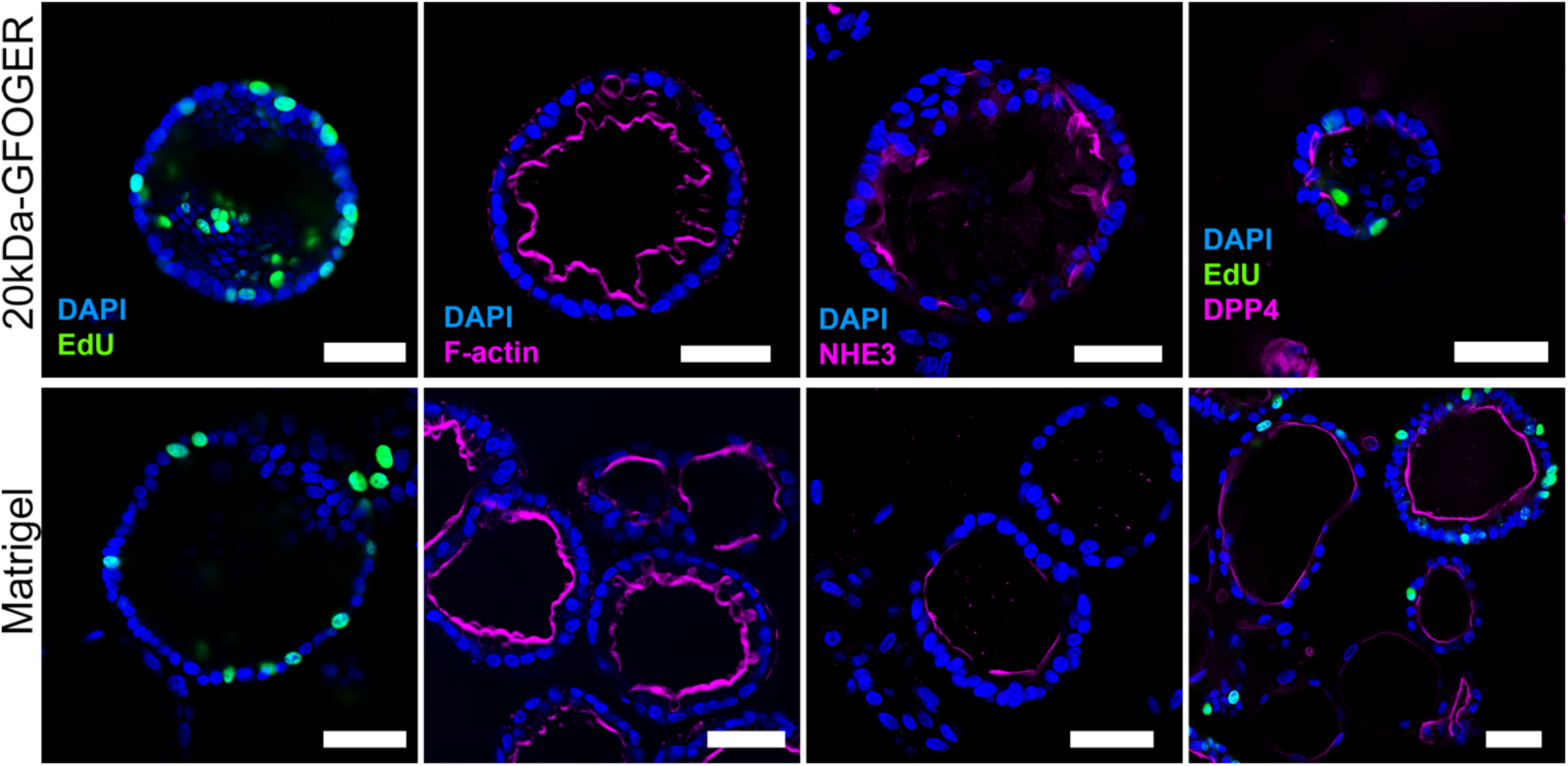
Human colon organoids in the synthetic ECM exhibit correct architecture and cellular composition. Immunostaining analysis of 10-day old organoids emerging in the synthetic matrix and Matrigel showing proliferative cells (Edu), and correct localization of apical markers (F-actin, NHE3 and DPP4).

**Extended data Fig 9.**
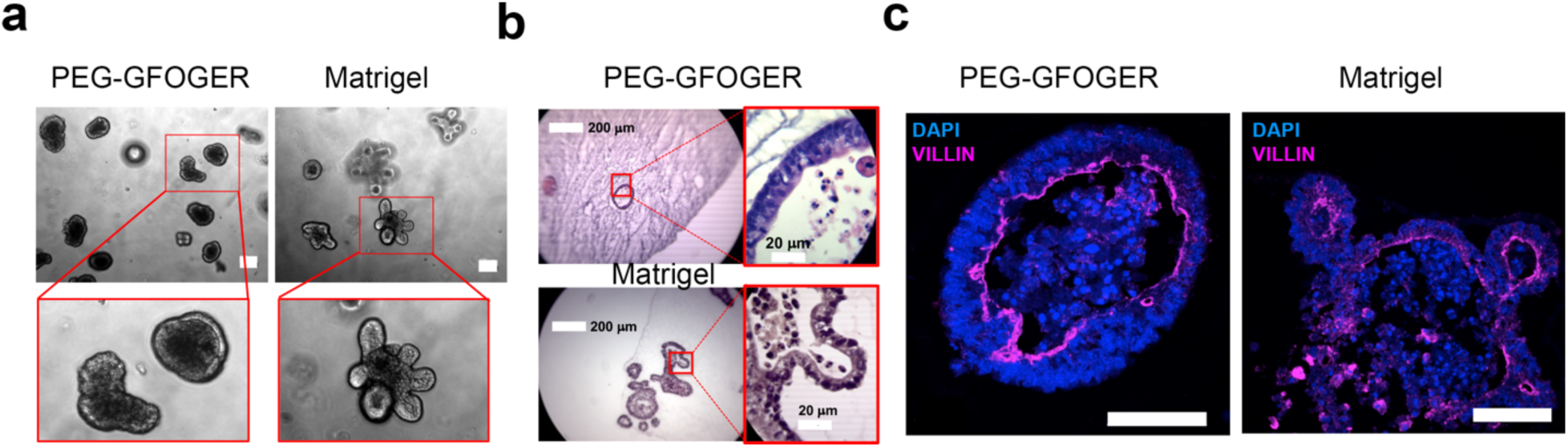
Mouse small intestinal organoids, in the synthetic ECM, undergo differentiation. **a**, Brightfield images of five-day old enteroids undergoing differentiation. Scale bar: 100 μm. **b,** Histological sections showing accumulation of apoptotic cells in the lumen upon differentiation. **c**, Immunostaining of differentiated organoids showing apical villin marker.

## Supplementary Data

**Table 1:** List of Taqman probes used for enteroid characterization.

| Name | Gene ID | Length (nt) |
| --- | --- | --- |
| GADPH | Hs02786624_g1 | 157 |
| LGR5 | Hs00969422_m1 | 61 |
| OLFM4 | Hs00197437_m1 | 85 |
| ASCL2 | Hs00270888_s1 | 101 |
| SMOC2 | Hs01591663_m1 | 80 |
| TROY | Hs00218634_m1 | 59 |
| BMI1 | Hs00995522_g1 | 153 |
| LRIG1 | Hs00394267_m1 | 96 |
| HOPX | Hs04188695_m1 | 56 |
| TERT | Hs00972650_m1 | 57 |
| WIP1 | Hs01013292_m1 | 65 |
| KRT19 | Hs00761767_s1 | 116 |
| KI67 | Hs00606991_m1 | 137 |
| HES1 | Hs00172878_m1 | 78 |
| DLL1 | Hs00194509_m1 | 74 |
| CHGA | Hs00900370_m1 | 67 |
| MMP7 | Hs01042796_m1 | 64 |
| ATOH1 | Hs00944192_s1 | 83 |
| TFF3 | Hs00902278_m1 | 64 |
| CCK | Hs00174937_m1 | 69 |
| LYZ | Hs00426232_m1 | 67 |

## Methods

### Data Reporting

The experiments were not randomized. The investigators were not blinded during experimental setup or evaluation of experimental outcomes. No statistical methods were used to determine sample size.

### Tissue processing

De-identified tissue biopsies were collected from unaffected intestinal regions of children and adult patients undergoing endoscopy for gastrointestinal complaints. Informed consent from adult donors or from a pediatric donors’ guardian along with developmentally appropriate assent donor were obtained at Boston Children’s Hospital. All methods were carried out in accordance with the Institutional Review Board of Boston Children’s Hospital (Protocol number IRB-P00000529). Tissue processing and initial human and mouse organoid culture were performed at the Harvard Digestive Disease Center, following published protocols^1, 2^. Organoids were grown in growth factor reduced (GFR) Matrigel droplets overlaid with expansion medium (EM) made with L-WRN conditioned medium (50% vol/vol, ATCC, CRL3276) and supplemented with Y-27632 (10 μM, TOCRIS)^1, 2^. Mouse colon organoids were cultured as previously reported^3^. Endometrial tissue collection was approved by the Partners Healthcare Institutional Review Board (Protocol number IRB-P001994). De-identified uterine tissue or pipelle biopsies were collected from consented adult donors (ages 18–45). Endometrial organoids were established using published protocols and cultured in endometrial organoid medium (EnOM)^5, 6^.

### Cell culture

Organoids were passaged every four days for human intestinal organoids, eight days for human endometrial organoids and three days for mouse organoids. For passaging, organoids were incubated in Cell Recovery Solution (CRS, Gibco) for 1h at 4C, followed by trypsin treatment to generate single cells. The cell suspension was inspected under the microscope to ensure the presence of dispersed single cells and if needed, filtered through a 30 μm cell strainer to remove cell clumps. Single cells were counted using a hemocytometer and seeded in GFR Matrigel (25 μL droplet) at a density of 1,000 cells/μL for human and 500 cells/μL for mouse organoids. After Matrigel polymerization, 600 μL of EM or EnOM was layered on top. Media was changed every two days.

### Organoid differentiation

Organoids in both Matrigel and the synthetic matrix were cultured in EM for six days (human) or four days (mouse) before switching to differentiation medium. Human and mouse organoids in Matrigel were differentiated as previously reported^1, 2^. Intestinal organoids in the synthetic matrix were differentiated using L-WRN conditioned medium (25% L-WRN) diluted in Adv DMEM/F12, HEPES (1 μM, Gibco), Pen/Strep (100 U/mL, 100 mg/mL, Invitrogen), N2 supplement (1X, Gibco), B27 supplement (1X, Gibco), human [Leu15]-gastrin I (10 nM, BACHEM), N-acetyl cysteine (1 mM, SIGMA) and Y-27632 (5 μM, TOCRIS). Mouse organoids in the synthetic ECM were differentiated as in Matrigel.

### PEG macromers and peptides

8-arm PEG macromers with vinyl sulfone (VS) terminal groups (40kDaPEG-VS and 20kDaPEG-VS) were obtained from JenKem Technology (Beijing). All peptides were custom synthesized and purified by Boston Open Labs (Cambridge, MA), GenScript (Piscataway, NJ), or CPC Scientific (Sunnyvale, CA). Peptides sequences and their relevant features are included in the Supplementary Table 1. “XL-VP,” a crosslinker containing a matrix metalloproteinase (MMP)-sensitive substrate, “XL-IA,” a crosslinker containing a MMP-sensitive substrate and a Sortase-sensitive recognition site (LPRTG), “XL,” a crosslinker containing only the Sortase-sensitive site, ‘‘RGDS,’’ a fibronectin (FN)-derived peptide containing the canonical RGD motif from the 10th FN type III domain, ‘‘PHSRN-K-RGD,’’ a FN-derived peptide containing the RGD motif and the PHSRN synergy site from the 9th FN type III repeat in a branched configuration; ‘‘GFOG**E**R,’’ a collagen I-derived peptide presented as a triple helix^8, 9^, “GFOG**D**R,” a collagen I-derived peptide with **E**➔**D** point mutation that reduces integrin binding^10^, “G11RGDS,” an extended ligand of RGDS with eleven Gly spacers, “CMPRGDS,” an extended and clustered RGDS ligand using a triple helical collagen mimetic peptide^11^, ‘‘FN-binder,’’ a peptide with affinity for FN, and ‘‘BM-binder,’’ a peptide with affinity for collagen IV (C-IV) and laminin (LMN)^12^. All peptides were reconstituted in acidic (pH 5.2) Milli-Q water (Millipore). The concentration of free thiols in all peptides was determined using Ellman’s reagent (Sigma).

### Synthesis of hydrogel precursors

All synthetic matrices were assembled at 5 % PEG (w/v) using 8-arm 40kDaPEG-VS or 8-arm 20kDaPEG-VS. The mixed 40/20kDaPEG synthetic matrix was made mixing 1:1 (v/v, 2.5% 8-arm 40kDaPEG-VS and 2.5% 8-arm 20kDaPEG-VS). All matrices were assembled stepwise via Michael-type addition reaction. First, the PEG-VS macromer was diluted with 10X PBS-HEPES solution (1 mM, pH 7.8); second, the matrix-binder peptides were added to the PEG reaction mixture and incubated for 30 minutes at RT; third, the integrin binder peptides (***RGDS***, ***PHSRN-K-RGD***, ***GFOGER***, ***G11RGDS***, ***CMPRGDS*** or **GFOGDR**) were added to the reaction mixture and allowed to react for an additional 30 min at RT. This sequential reaction created a PEG-functionalized mixture ‘‘fPEG-VS’’ that was used to resuspend the cells prior to matrix gelation (see cell encapsulation below). The nominal concentration of the matrix-binders in all matrices was 0.25 mM each, whereas the integrin binder peptides were at 1.5 mM (unless otherwise noted in the figure legends). For most experiments the fPEG-VS solution was crosslinked at 50% crosslinking density (unless otherwise noted in the figure legends).

### Rheological characterization

After adding the XL-IA crosslinker (50%), 20 μL of each matrix mixture was loaded into a 1 mL syringe that had the tip cut off at the 0.1 mL mark. The matrix was allowed to gel at 37 °C for 20 min and then moved to a 24 well plate that contained 400 μL of 1X PBS. The plate was incubated for 24 hours in a humidified incubator at 37 °C, 95% air, and 5% CO_2_, to allow for equilibrium swelling to occur prior to rheological characterization. This procedure created matrices discs of 1–1.4 mm in thickness. The discs were sandwiched between an 8 mm sandblasted parallel plate and sandblasted base. The shear modulus was determined by performing small-strain oscillatory shear measurements on an Anton Parr MCR 302 instrument. The mechanical response was recorded by performing frequency sweep measurements (0.1–10 Hz) in a constant strain mode (0.05), at 37 °C. The shear modulus (G’) is reported as a measure of matrix mechanical properties.

### Equilibrium swelling

The matrices were prepared at 50% crosslinking density (unless otherwise noted in the figure legends) as described above. After adding the XL-VP crosslinker, three 30-μL droplets were loaded onto three 18-mm circular glass micro-coverslips.

The mass of the coverslip (mCs) were determined prior to the addition of the gel mixture to calculate the percentage of swelling (see below). The gel mixture/coverslips were placed inside of a 12-well plate and allowed to gel for 20 min in a humidified incubator at 37 °C, 95% air, and 5% CO_2_. At the end of the gelation, each coverslip was weighed again and the mass recorded as “pre-swelled matrix mass (mpSM)”. One mL of 1X PBS was loaded onto each matrix droplet and returned to the incubator for 24 hours to allow the matrix to reach equilibrium swelling. After 24 hours, the PBS was removed, and the matrices were washed twice with 1 mL of dH_2_O. The water was removed completely before the matrix/coverslips were weighed again. This was recorded as “swelled matrix mass (mSM)”. Finally, the matrix/coverslips were placed in a 60 °C oven overnight to determine the mass of the dry matrix. To calculate the percentage of swelling we sued the following formula ((mpSM-mCs)/(mSM-mCs))*100. The pore size (ξ) was calculated according to Flory-Rehner equations and derived formulas described previously ^13^. In experiments to determine the effect of the crosslinking density on swelling and pore size, the synthetic matrices were made with 35, 40, 45, 50, 55, and 60% XL-VP crosslinker density. The percentage of swelling and pore size were calculated as above.

### Cell encapsulation

Four-day old organoids grown in Matrigel were collected and processed as above to generate single cells. The cell suspension was inspected under the microscope to ensure the presence of dispersed single cells and if needed, filtered through a 30 μm cell strainer to remove cell clumps. Single cells were counted using a hemocytometer then resuspended in the matrix precursor solutions (fPEG-VS) prior to the addition of the crosslinker and Y-27632 (10 μM). In parallel, single cells were resuspended in Matrigel that served as experimental control during matrix evaluation. In both cases, cells were encapsulated at a density of 500 cells/μL of matrix. Three μL (1,500 cells) of the matrices were loaded into a Nunc MicroWell 96-well optical-Bottom plate and allowed to polymerase for 20 min in a humidified incubator at 37 °C, 95% air, and 5% CO_2_. After gelation, 100 μL of EM or EnOM was loaded into each well. Media was changed every two days. Eight-day old endometrial organoids were used to generate single cells. The encapsulation process in Matrigel or the synthetic ECM, for endometrial organoids, was performed similar to intestinal organoids.

### Intestinal and endometrial enteroid/organoid diameter

The diameter of six-day old enteroids and eight-day old endometrial organoids were determined using a deep learning based algorithm as described previously^14^. 4X brightfield (BF) images were captured using an EVOS M500 microscope (Invitrogen).

### Enteroid formation efficiency

Single cells were encapsulated as above and cultured in EM for four days (Matrigel) or six days (synthetic matrix) before imaging at 10X magnification using an EVOS M500 microscope. Sixty BF images, spanning the entire thickness of the matrix, were collected from the center of the droplet. This is approximately 1/3 of the total droplet volume. The image z-stacks were processed in Fiji using the time lapse Gaussian-based stacker focuser plugin to create a single image with all enteroids in focus. The number of enteroids with a clear lumen and the number of single cells were manually counted using the cell counter feature in Fiji^15^. We also observed and counted few cell clumps that did not show a clear lumen but were bigger than single cells. Enteroid formation efficiency was calculated as the percentage of enteroids with a clear lumen relative to the total number of enteroids, single cells, and cell clumps, counted in each droplet.

### Live/dead imaging

Single cells were encapsulated as before for six days (intestinal) or eight days (endometrial) before the addition of Calcein AM (2 mM) and Ethidium homodimer-1 (2 mM) for 20 minutes. Images were captured using either a ZEISS confocal Laser Scanning Microscope (LSM 880) equipped with temperature (37 °C), humidity, and CO_2_ (5%) controls orr an EVOS M500 (Invitrogen) microscope (no incubation). A 1.6 mm by 1.6 mm area and ∼800 μm thick section of either Matrigel or the synthetic ECM were imaged with the confocal. With the EVOS, we captured the center of the droplet, which is ∼ 1/3 of the total matrix area. The final images were processed using the ZEN blue ZEISS companion software or Fiji^15^. For time-course live/dead imaging analysis, 60 z-stacks images were taken the day of seeding (day 0) then every two days for up to ten days, using the EVOS M500 microscope and processed as above.

### Continues live imaging

Single cells were encapsulated and plated on 96-well optical plates as described before, then cultured in OEM. Bright field images were captured using a ZEISS confocal Laser Scanning Microscope (LSM 880) equipped with a wide-field camera and temperature (37 °C), humidity, and CO_2_ (5%) controls. A single plane of four-day old enteroids were imaged every 5 minutes for 48 hours. The final videos were prepared using Fiji using frame interpolation to smooth the video^15^. To capture individual cells forming enteroids, the entire thickness of the matrices was imaged the day after encapsulation and then the everyday for up to six days. The stack of images was processed in Fiji as described in enteroid formation efficiency ^15^.

### Cell proliferation

To determine if cells from enteroids in the synthetic ECM retain their proliferative capacity, we collected six-day old enteroids from within the synthetic ECM using SrtA treatment^16^, then digested them with trypsin to generate a single cell suspension. Cells (500 cells/μl, 5 μl droplets, 48 droplets total) were embedded in Matrigel or the synthetic ECM and cultured in EM for six days. After six days, the matrix droplets were pooled and processed as follows; enteroids in Matrigel were released using CRS whereas enteroids in the synthetic ECM were released using SrtA treatment. Enteroids in both conditions were then digested with trypsin to get the total number of cells recovered from the pooled enteroids, from each matrix condition. The fold increase in cell number was calculated as the ratio of total number of cells obtained from enteroids after six days of culture divided by the total number of cells used at the beginning of the experiment. The cells recovered from enteroids in Matrigel were discarded. The cells recovered from enteroids in the synthetic ECM were used to set up a new experiment (first passage, 500 cells/μl, 5 μl droplets, 48 droplets total) in new synthetic ECM or Matrigel. At the end of six days, the fold increase in cell number was determined as before. This process was repeated three consecutive passages using two human duodenal donors. In parallel experiments, using the same pool of cells, we measured the enteroid diameter of the three consecutive passagesas previously described^14^.

### Quantitative real time PCR (qPCR)

Four-day old enteroids grown in Matrigel were used to generate a single-cell suspension as described above. Single cells (500 cells/μL) were encapsulated in Matrigel or the synthetic ECM, then cultured in EM. After six days, intact enteroids were released from Matrigel and the synthetic ECM, as described before. Intact enteroids were pelleted, resuspended in TRIzol reagent (ThermoFisher Scientific, 15596026), and then stored at −80 °C until processing. From the initial cell suspension, we reserved 500,000 cells at −80 °C in TRIzol. This single-cell population was used to determine the gene expression of the “initial cell population”. RNA was extracted from enteroids (or cells) using the Directzol RNA Mini-Prep kit (Zymo Research) per the manufacturer’s protocols with the inclusion of an on-column DNase step using the PureLink DNase Set (Thermofisher Scientific, 12185010). cDNA was synthesized from ∼ 1 μg of total RNA using the High-Capacity RNA-to-cDNA Kit (Thermofisher Scientific, 4387406) per manufacturer’s protocols. TaqMan Fast Advanced Master Mix (Thermofisher Scientific, 4444557) was used in congruence with the cell-specific probes for qPCR described in the supplementary Table 2. Gene expression was determined using the StepOnePlus real-time PCR system (Applied Biosystems) and calculated using the ΔΔCt method in GraphPad Prism. Gene expression was first normalized using the housekeeping GAPDH gene in each sample, then the relative fold change in gene expression, in Matrigel or the synthetic ECM, was calculated against the gene expression of the “initial cell population” that was set to 1. The experiment was repeated three times with two duodenal donors.

### Histological processing and immunostaining

Mouse intestinal organoids were fixed in while still in the gel (3D), then paraffin-embedded and sectioned in the Histology Center at the Koch Institute at MIT. 5-micron tissue sections were hematoxylin and eosin stained using standard procedures.

### Immunostaining and EdU labeling

Organoids and enteroids were processed in two formats: in 3D (embedded within the synthetic ECM or Matrigel) and in suspension (free floating) after being released from the matrices. Six-day old enteroids in 3D were treated with EdU (5-ethynyl-2’- deoxyuridine, 20 μM) for 3 hr (Alexa Flour 488 Click-it EDU, Thermo Fisher) prior fixation overnight with formalin (10%, VWR) at 4 °C. Eight-day old endometrial organoids in 3D were treated with EdU for 6 hr prior to overnight fixation. Eight-day old colon organoids were treated with EdU for 6 hr, then released from the matrices and fixed as free-floating organoids for 30 min at RT. After fixation, organoids and enteroids in 3D were permeabilized with 0.1% triton X-100 in PBS overnight followed by blocking (4% BSA/0.5% Tween 20 in 1XPBS or 4% Donkey serum/0.5% Tween 20 in 1X PBS) overnight at 4°C with 200 rpm shaking. Free-floating organoids were permeabilized for 1h at RT incubated in a tube rotator set at 30 rpm, followed by 3 hr of blocking. To identify proliferative cells, the Alexa Flour 488 was click-reacted according to the manufacturer instruction. Immunostaining for cell-specific markers was done using the following primary antibodies, diluted in blocking solution; goat anti-E-cadherin (R&D, AF748, 1:200), goat anti-hDPPIV/CD26 (Invitrogen, AF1180, 1:400), rabbit anti-Col IV (Abcam, Ab6586, 1:200), rabbit anti-Lysozyme (Dako, A0099, 1:200), rabbit anti-LMN (Abcam, Ab11575, 1:200), rabbit anti-Ki67, (Abcam, Ab15580, 1:200 or ab16667, 1:100), rabbit anti-NHE3/SCL9A3 (Novus, NBP-82574), mouse anti-Villin (Santa Cruz, SC-58897, 1:50), mouse anti-CD44-v6 (Abcam, Ab78960, 1:200), mouse anti-Muc2 (Santa Cruz, SC515032 1:200), mouse anti-EpCAM (Abcam, ab7504 1:200). Samples in 3D were incubated for two days in primary antibodies at 4°C and 200 rpm shaking. Free-floating samples were incubated overnight in primary antibodies at 4°C in the tube rotator set at 30 rpm. The secondary antibodies and dilutions used were; donkey anti-goat Alexa Fl 568 (ThermoFisher, A11057, 1:200), donkey anti-rabbit Alexa Fl 568 (ThermoFisher, A10042, 1:200), and donkey anti-mouse Alexa Fl 568 (ThermoFisher, A10037, 1:200). Nuclear staining was done using DAPI (1 mg/mL, 1:2,000). F-actin staining was done using either Alexa Fl 488 phalloidin (ThermoFisher, A12379, 1:200) or Alexa Fl 568 (ThermoFisher, A12380, 1:200). DAPI and f-actin staining was done in combination with the secondary antibody. Samples in 3D were incubated in secondary antibodies for two days at °C and 200 rpm shaking. At the end of the incubation, samples in 3D were washed five times (10 minutes each, 200 rpm at RT) prior to the addition of CytoVista reagent (Thermofisher) to match the refraction index and allow imaging in 3D. Free-floating samples were incubated overnight in secondary antibodies at 4°C in the tube rotator set at 30 rpm, followed by five washes (10 min each, at RT and 30 rpm). The organoids were gently pelleted and resuspended in prolong gold antifade reagent (ThermoFisher, P36935), then transferred onto a glass slide modified with a 20 mm secure seal spacer (0.12 mm deep, ThermoFisher, S24736). 5-micron tissue sections were incubated in mouse anti-Villin (Santa Cruz, SC-58897, 1:50) overnight, followed by donkey anti-mouse Alexa Fl 568 (ThermoFisher, A10037, 1:200) and DAPI staining. Images were captured using a ZEISS confocal Laser Scanning Microscope (LSM 880).

### Statistical analysis and sample information

Statistical significance between experimental treatments were determiend as follows: Enteroid diameters were analyzed using one-way ANOVA and Kruskal-Wallis multiple comparison of the mean ranks. Storage modulus data analyzed using one-way ANOVA and Sidak’s multiple comparison whereas swelling and pore size data was analyzed using Kruskal-Wallis multiple comparison of the mean ranks. Enteroid formation was analyzed using either one-way ANOVA and Holm-Sidak’s multiple comparison of the mean or two tailed Mann-Whitney tests when comparing two groups. qPCR data were analyzed using multiple t tests and the Holm-Sidak method. All statistical analyses were performed in the GraphPad Prism 8.0 software.

## Data availability

Source Data for Figs. 1–6 are provided with the paper. All additional relevant data are available upon request from the corresponding author.

## Acknowledgements

We thank the Harvard Digestive Disease Center for establishing the original intestinal cultures from biopsies. Professor Thaddeus Stappenbeck from Washington University School of Medicine in St. Louis for providing the mouse colon organoids and the L-WRN cell line to produce the conditioned medium. We thank the Koch Institute histology and microscopy centers at MIT for helping establish protocols for organoid processing. We thank Alexander Brown (MIT) for helpful discussions of the manuscript, Dr. Katherine Bezold Lamm (MIT) for help collect some samples for qPCR analysis and for her helpful comments on the manuscript and Derrick Marsall (MIT) for help in the rheological characterization. We thank the MIT UROP office for financial support to A. L. and G.C. This work was supported by funding from NIH Research Grant No. R01EB021908 and by financial support from DARPA and Boehringer Ingelheim through the SHINE program.

## Author Contributions

V.H.G. and L.G.G conceived the study, designed experiments and wrote the manuscript. V.H.G. cultured organoids and performed the experiments with the help of A. L, G. C. and M.G. All were involved in data collection whereas V.H.G was involved in processing and analyzeing all data. A.L. performed matrix characterization. T.K. conceived and developed the algorithm to measure the enteroids, help processing data and edited the manuscript. J.G.N. collected, processed and established human endometrial organoids and performed immunostaining of endometrial organoids. R.C. edited the manuscript. D.B. supervised tissue collection to establish initial organoid culture, edited the manuscript.

---

**Supplementary Table 1.**
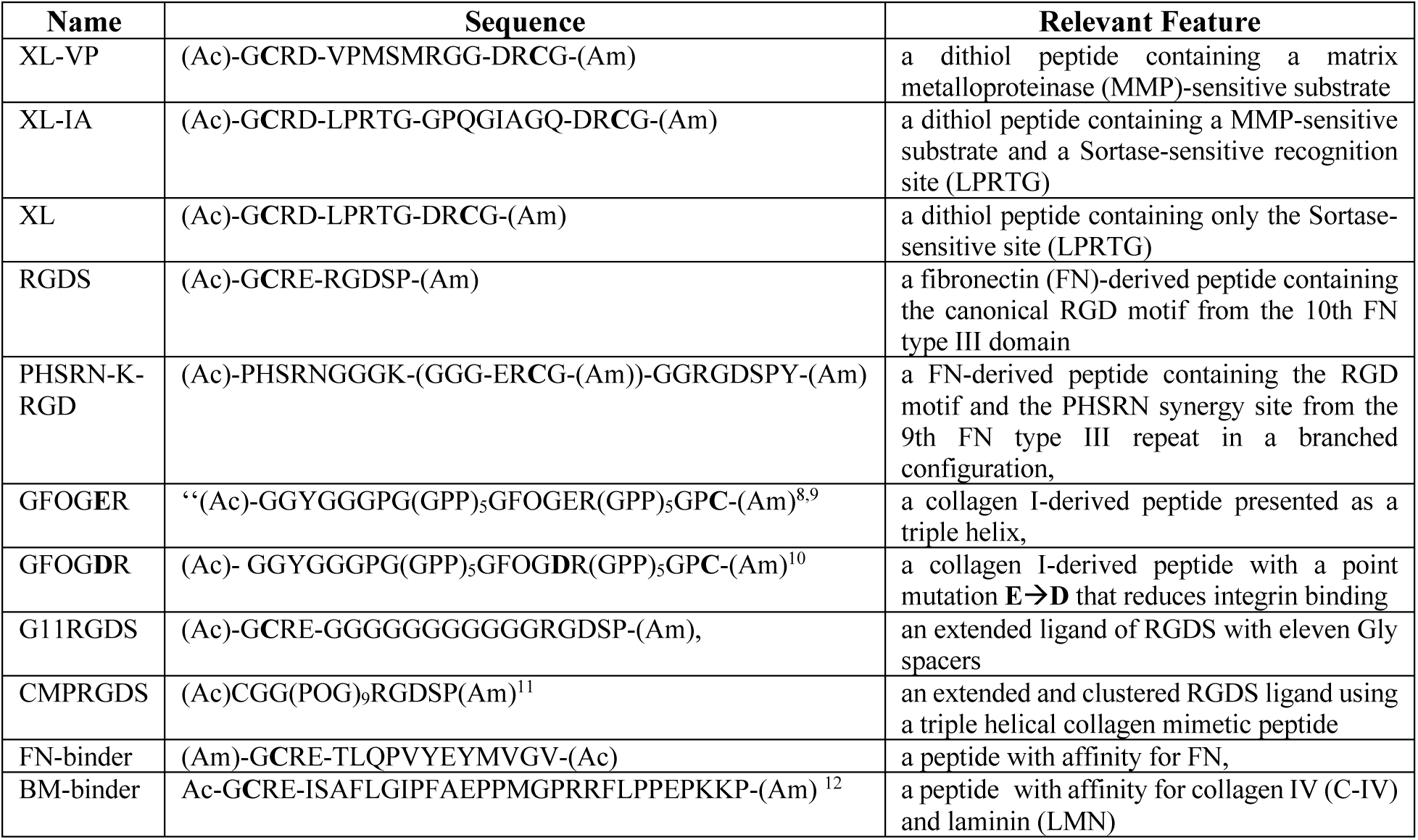
Peptide sequences and relevant features.

